# Optogenetic control of pheromone gradients and mating behavior in budding yeast

**DOI:** 10.1101/2024.02.06.578657

**Authors:** Alvaro Banderas, Maud Hofmann, Céline Cordier, Matthias Le Bec, M Carolina Elizondo-Cantú, Lionel Chiron, Sylvain Pouzet, Yotam Lifschytz, Wencheng Ji, Ariel Amir, Vittore Scolari, Pascal Hersen

## Abstract

During mating in budding yeast, cells use pheromones to locate each other and fuse. This model system has shaped our current understanding of signal transduction and cell polarization in response to extracellular signals. The cell-population produced extracellular signal landscapes themselves are however less well understood, yet crucial for functionally testing quantitative models of cell polarization and for controlling cell behavior through bioengineering approaches. Here we engineered optogenetic control of pheromone landscapes in mating populations of budding yeast, hijacking the mating-pheromone pathway to achieve spatial control of growth, cell morphology, cell-cell fusion, and distance-dependent gene expression in response to light. Using our tool, we were able to spatially control and shape pheromone gradients, allowing the use of a biophysical model to infer the properties of large-scale gradients generated by mating populations in a single, quantitative experimental setup, predicting that the shape of such gradients depends quantitatively on population parameters. Spatial optogenetic control of diffusible signals and their degradation provides a controllable signaling environment for engineering artificial communication and cell-fate systems in gel-embedded cell populations without the need for physical manipulation.

## 1. Introduction

Cell-cell communication through extracellular signals enables the spatiotemporal coordination of cell behavior across various scales, from single cells to tissues. The interplay between signal production, diffusion, and reaction (*e.g.*, internalization, adsorption, and degradation) shapes the concentration landscapes that cells experience and ultimately determines a cell’s responses. A classic example is the establishment of positional information in the *Drosophila melanogaster* syncytial embryo, in which an exponentially decaying extranuclear *bicoid* morphogen gradient forms as a result of the specific location of emitter nuclei on the anterior region, free diffusion of the signal, and homogenous degradation of the signal throughout the embryo^1,2^. This smooth gradient is “read” by nuclei to switch on specific differentiation programs in sharply defined spatial locations.

In contrast to multicellular developmental systems where cells follow a predefined program from a well-defined initial condition, communication does not necessarily occur within closed spaces for free-living microbes. Populations of single cells often have a random spatial distribution, expand freely, explore an unknown environment, and create an ecosystem—the size and dynamics of which are the result of cellular metabolic interactions and cell-cell communication. As in development, signal landscapes depend on reaction-diffusion dynamics; however, population-level parameters such as cell density, population composition, and the spatial distribution of emitter and receiver cells are inherently variable in the context of single-cell populations. Thus, the way cells extract information and take appropriate decisions in such an uncertain environment is a very different biological problem than what is observed in embryology.

In the well-known example of mating through pheromones in the budding yeast *Saccharomyces cerevisiae*, information gathered by cells is usually considered positional, *i.e.*, gradients formed will attract cells to polarize towards the position of the source. It is generally accepted that, in nature, such positional information is used by cells only over very short (“touching”) distances from partners^3^. This is in agreement with ecological lifestyle of *S. cerevisiae*, where mating-type switching^4^ and intra-ascus mating^5^, where mating cells are placed in close contact to each other, ensure fast return to the diploid lifestyle.

The molecular mechanism underlying gradient tracking involves polarisome movement (with both deterministic^6,7^ and stochastic stages^8,9^) in response to pheromone-producing adjacent cells. During mass mating reactions, cells display mostly such “short-range” chemotropism (**Fig. 1A**), where growth and cell-body extension beyond one cell length does not occur prior to fusion. In contrast, cells in microfluidic devices, subject to artificial gradients, display what we can call long-range chemotropism (**Fig. 1A**), where the whole cell body extends, greatly increasing the length, cell surface and volume of cells. Additionally, filamentous growth, where cells extend and bud in the direction of gradients, has been observed in pheromone disc assays^10^.

**Figure 1.**
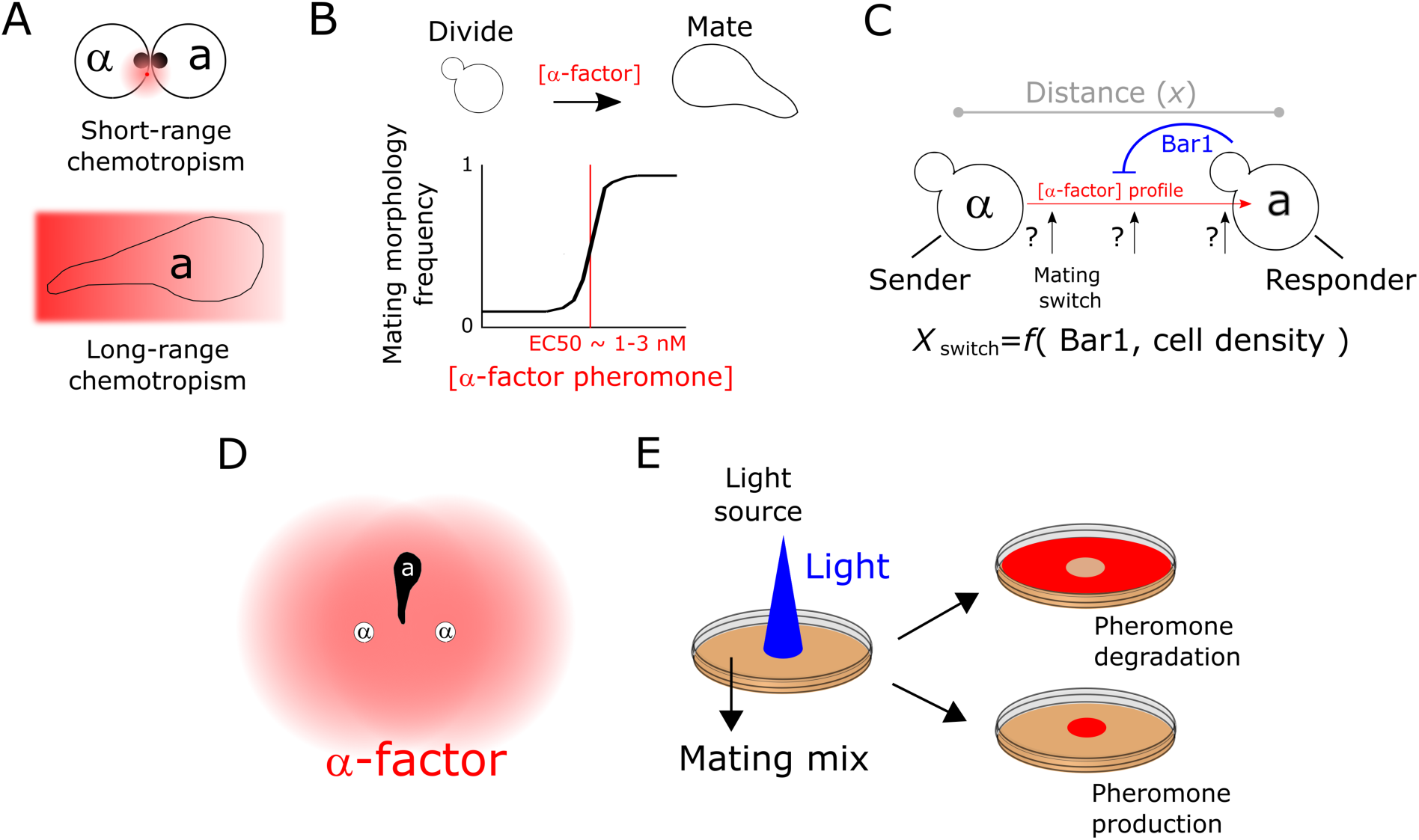
Using optogenetics to interrogate the properties of cell-generated signaling gradients. **A.** Two types of chemotropism have been described in the literature (see text), short range (top) with no cell-elongation involved and frequently observed in mating reactions (black dot represents the polarisome and pheromones—pink halo—are secreted locally), and long-range (displaying cell elongation), seen in artificial pheromone gradients. **B.** The mating switch. Yeast cells switch from budding to mating morphologies when α-factor is present (top). This dose-response is sharp and occurs in the nM range, close to the receptor Kd across strains. **C.** When cells with opposite mating types are located at a distance, the generated α-factor gradient (red arrow, indicating the direction of decay) has a strength (amplitude and steepness) that defines where the concentration is sufficient for switching-on mating behavior. The profile of the α-factor pheromone not only depends on the local density of emitters contributing to the gradient, but also on the presence of the extracellular α-factor peptidase Bar1, and therefore on the density of *MAT***a** cells. When Bar1 is absent or low, the position of the mating switch is expected to happen at a distance. In contrast, if Bar1 is high, the mating switch is displaced towards the source vicinity. **D**. Due to low Bar1 mediated degradation of α-factor when *MAT***a** populations have low density, the α-factor gradients are expected to overlap and provide sufficient intensities to trigger mating behavior at a distance, but at the cost of reduced precision due to “confusing” local maxima. **E.** Optogenetic control of α-factor and Bar1 emission allows spatial control of these two opposing activities. In the example shown, a Petri dish containing mating mixes is illuminated in specific regions allowing us to stimulate production or degradation of the pheromone (red color), depending on which genetic determinant is controlled.

Chemotropic behaviors suggest a role for cell elongation in directed mate-finding. Such behavior is potentially relevant for sparse populations generated after spore dispersal events in wild yeast^11,12^, where mating type switching is not the rule. Indeed, wild isolates of *S. cerevisiae* display large differences in their propensity to mate or to enter the cell cycle and thus spores germinating in the wild may often form microcolonies before engaging in mating^11^. Importantly, as in bacterial quorum sensing^13^, “demographic” information has also been shown to be a relevant cue in sparse mating populations^3,14,15^. These examples highlight the idea that mating responses are not restricted only to close-contact interactions in nature, and that population-parameters could be therefore relevant in both natural and synthetic ecosystems.

Mating in yeast is governed by the mating-pheromone pathway,^16,17^ which relies on the secretion of pheromones to signal the presence of different mating types (*MAT***a** and *MAT*α). The *MAT***a** and *MAT*α mating-types exchange pheromones to trigger activation of their mating pathway, orchestrating a dedicated transcriptional program, cell cycle arrest, and morphological changes that culminate with cell and diploid formation. Distinct branches of the pheromone pathway control transcriptional (Ste12 branch) and arrest/morphology behaviors (Far1 branch). Dose-response experiments measuring the responses of *MAT***a** cells to the purified pheromone α-factor in bulk showed that, while the transcriptional branch of the pathway displays a Michaelian (linear at low doses) input-output relation^18^, cells transition sharply—following a Hill-type sigmoidal response— from proliferating to growth-arrested elongated phenotypes (called “shmoos”) characterized by their mating protrusions ^19,20^. This “mating switch” **(Fig. 1B)** was notably observed in artificial gradients of α-factor^21,22^ generated using microfluidic devices. In this setting, cell cycle-arrested morphologies were observed over a narrow range of concentrations (1‒5 nM) of α-factor. Cells only displayed chemotropic behavior, elongating their cell body up the α-factor gradient, within this concentration range. However, importantly, it remains unknown how the amplitude and steepness of artificially generated gradients in the context of microfluidic devices compare to those within heterogeneous, large populations of mating cells. Indeed, elongation with full accuracy of elongated-cell chemotropism in microfluidic chambers is generally observed at high gradient and amplitude values^9,19,21–24^ (hundredths of pM/µm). Thus, the question remains: under which conditions mating populations produce such specific gradients such that cells undergo useful long-range chemotropism?

In addition to responding to α-factor, *MAT***a** cells secrete the Bar1 protein which degrades α-factor extracellularly. Thus, the pheromone landscape^3^ that *MAT***a** cells experience is not only determined by spatiotemporal diffusion of the α-factor pheromone produced by *MAT*α cells, but also by the concentration of Bar1 they produce themselves. For a non-motile organism such as *S. cerevisiae*, evolution would be expected to produce systems where the concentrations of α-factor that trigger mating are found only in the vicinity of emitter cells, so that cells do not have mating responses unless they are very close to a partner. Bar1 has been deemed responsible to exert such a function^25^, by increasing the degradation rate of α-factor and confining its activity close to the producer cells, consequently shortening the length-scale at which mating sensing occurs to only a few micrometers (a fraction of cell size). However, in sparse, growing mating reactions, the concentration of Bar1 is dependent on the density of *MAT***a** cells^14^; therefore, the levels of Bar1 may not always be sufficient to degrade the α-factor produced by cells of the opposing mating type. This implies that the mating-switch distance can be, in theory, extended further away from the source position (**Fig. 1C**). Consequently, the superimposition of gradients produced by multiple pheromone sources represents a potential confounding factor (**Fig. 1D**): partners located at a distance from source cells cannot decide with which partner to mate^26^. Indeed, in mating discrimination assays^26^, *bar1Δ MAT***a** can discriminate real emitters from dummy (that do not produce pheromone) emitters, but only if real emitters are in the minority. This suggests that in the absence of degradation, pheromone halos from other cells can be a confounding factor for receiver cells. In summary, it remains unclear how cells extract relevant positional information from physiological pheromone concentration landscapes^3^.

Several theories have attributed various possible functions to Bar1 in mating populations. Three of these theories depend on high local concentrations of Bar1 in receiver cells, acting as a localized sinks of α-factor: (i) reducing the size of the pheromone halo around emitters (at the expense of gradient strength) to correctly align to gradients in the presence of multiple sources^27^, (ii) gradient disentanglement from two sources in close-proximity (intra-ascus) interactions,^28^ and (iii) mutual-avoidance in *MAT***a** cells^29^. Despite these functions requiring direct interactions of Bar1 with the cell wall, a large fraction of Bar1 is considered to be widely diffusible.

Understanding the spatial properties of α-factor-mediated communication between mating types requires characterizing the properties of physiological α-factor gradients. Although the shape of gradients can be inferred using numerical simulations of cells randomly arranged in space^30^, the inability to experimentally control α-factor production locally and the high response variability among receivers preclude the construction of predictive models. Here, we address this problem using the power of optogenetics (**Fig. 1E**). We developed a method for the spatial control of α-factor’s production and degradation in *Saccharomyces cerevisiae* and obtained a quantitative description of extracellular α-factor gradients under physiological conditions. Combined with a diffusion-reaction model, we inferred quantitative parameters that describe the mating communication system, providing direct quantitative support to indirectly inferred pheromone gradient and response properties in a single assay. In synthesis, we provide proof of principle for the optogenetic quantitative control of extracellular signal gradients within microbial cell populations.

## 2. Results

### 2.1 Engineered optogenetic induction of physiological range α-factor levels allows spatial control of mating

We first aimed at engineering light-inducible strains that can produce physiological levels of pheromone, by improving on a previous strain design^31^. For this we first engineered wild-type *MAT*α yeast cells using a CRISPR-based promoter replacement strategy (**see Methods**) to replace the promoter of the two α-factor pheromone-coding genes (MF-α1 and MF-α2) with the P_C120_ promoter, the activity of which is induced by the light-sensitive transcriptional activator EL222^32^. Additionally, to increase α-factor production and recover wild-type levels, a second copy of the P_C120_-MFα1 construct was inserted at an unrelated locus (**Fig. 2A, Figure S1, see Methods**) to yield the “opto-α” strain used in this study. To quantify the levels of α-factor produced by the opto-α strain, we used a *MAT***a** reporter strain that harbors a super-folder green fluorescent protein (GFP) downstream of the P_FUS1_ pheromone-responsive promoter^14^ (**Fig. 2B**). This reporter strain harbors the native untranslated terminal region (UTR) of FUS1 and carries a ubiquitination sequence that targets super-folder GFP for faster degradation, thus providing a good estimate of the abundance of the native transcript^14^. We stimulated mixes of opto-α and reporter strains with light using a custom-made light-plate apparatus^33^ and quantified the levels of the α-factor pheromone (relative to the wild-type MATα strains) using two reporter strains that either carry or lack the Bar1 protease (**Fig. 2B**).

**Figure 2.**
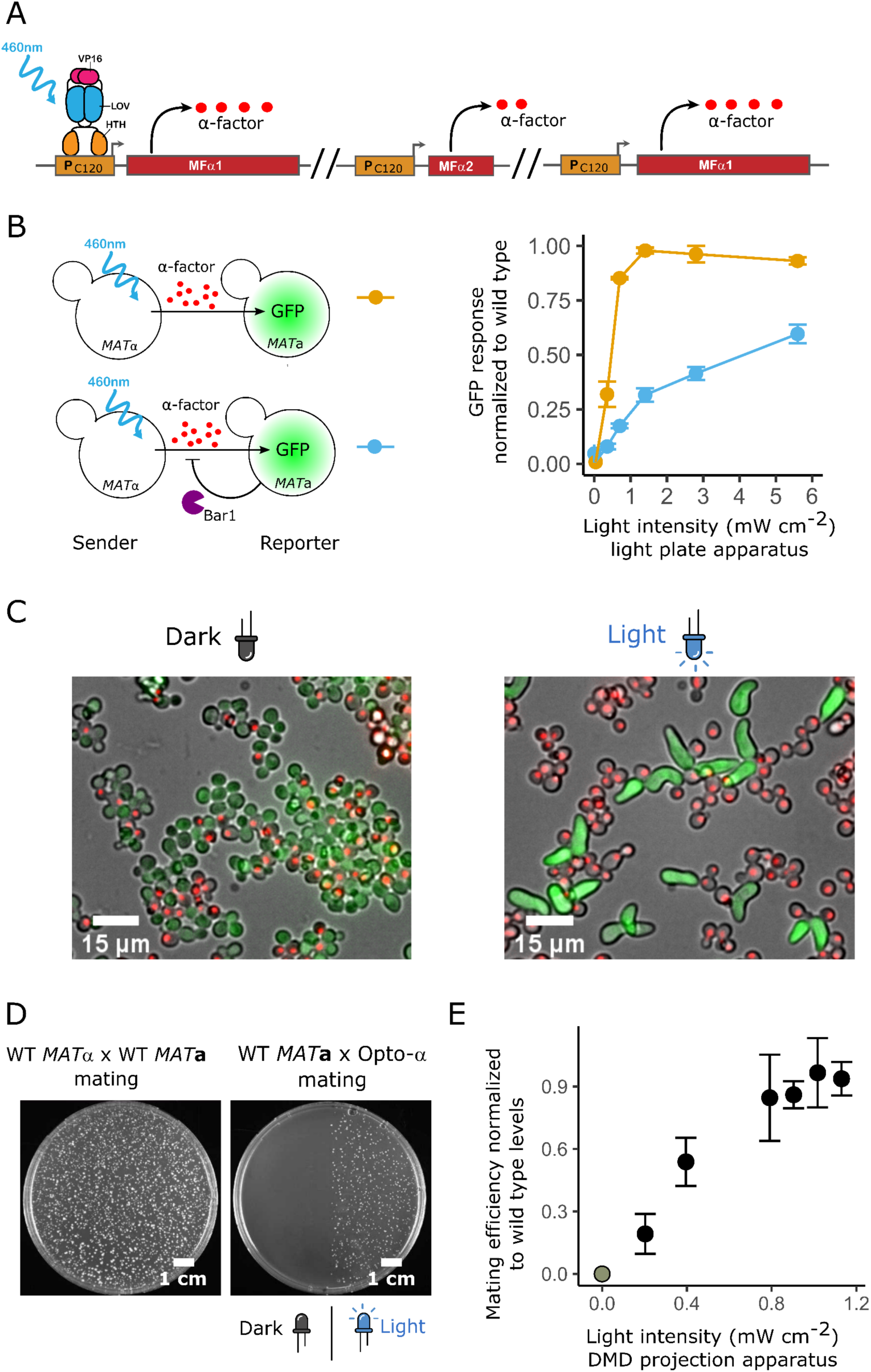
Optogenetic control of extracellular α-factor production allows tight spatial control of mating. **A.** The light-responsive α-factor-producing “opto-α” strain was engineered by replacing the native promoters of the two pheromone-coding genes in *MAT*α (MFα1 and MFα2) with the P_C120_ promoter, which is responsive to the light-activated LOV domain EL222 transcription factor, and by later duplicating the MFα1 (which produces the highest amount of pheromone of the two genes) light-responsive locus. **B.** The activity of the P_FUS1_ pheromone-responsive promoter in *MAT***a** receivers in stirred mating reactions as a function of light intensity. In the assay, the opto-α strain is co-incubated either with *bar1*Δ (top left) or a wild-type *MAT***a** (bottom left) P_FUS1_ GFP reporter strain. After 2 h, the mixes are assessed by flow cytometry. As a control, the reporters are incubated with wild-type *MAT*α, and the baseline-subtracted relative response is calculated (right). Error bars are the SD of n=3 biological replicates. **C.** Agarose-embedded *MAT***a** bar1Δ reporter strains (cells carrying cytoplasmic GFP) co-incubated with opto-α cells (carrying a red nuclear signal expressed by a mApple fluorescent protein) for 5 h under non-illuminated (left) and maximally illuminated (5.8 mW/cm², right) conditions. **D, E.** Quantification of mating efficiency through light-controlled pheromone production. Light induces cell-cell fusion in the optogenetic mating assay measured as the appearance of diploid colonies only in illuminated regions (D), which was quantified at different light-doses (E). Efficiency is defined as relative to the activity of the wild-type *MAT*α strain (C). Error bars are the SD of n=3 biological replicates.

In both cases, the levels of α-factor were light-intensity dependent. In the absence of Bar1, pheromone levels reached the induction levels achieved by the wild type. Under the more stringent conditions where receiver cells express Bar1, the opto-α strain reached up to ∼60% of the wild-type response. As (i) other strains and conditions resulted in lower α-factor production levels (**Fig. S1,S2**), (ii) sample treatment of optogenetic strains puts them at disadvantage in terms of pheromone production as, comparatively, wild-type cells constantly produce Bar1 during pre-illumination processing (see methods), and (iii) in general mating is very robust to various pheromone levels^34^, we decided to focus on the opto-α strain and decided to score its wildtype physiological pheromone production by checking if the levels of pheromone produced can rescue wild-type physiological behavior. For this, we tested the capacity of *MAT***a**-*bar1*Δ reporter cells to develop mating protrusions in response to light-induced α-factor production by opto-α cells. With sufficient intensity of light and, like with wild-type *MAT*α emitters, we observed formation of protrusions by *MAT***a** *bar1*Δ cells in gel-embedded mixes (**Fig. 2C, see Methods**). Finally, we assayed diploid formation in opto-α/*MAT***a** mixes as a measure of the final physiological output of the mating pathway (**Fig. 2 D,E**). For this, we developed an “optogenetic mating assay” (**Fig. S3, see Methods**) based on three features. First, we used the top-agar technique to embed mating mixtures (as used in classical growth arrest assays), which provides a homogeneous layer where cells are located sparsely and grow as microcolonies. Second, we employed a digital micromirror device (DMD)^35^ to expose the cells to spatial light patterns projected into the agar (**See Methods**); and finally, auxotrophic markers were used to ensure that only diploid colonies can grow, allowing us to have a measure of light-induced mating efficiency. Our results show that pheromone production is indeed sufficient to induce cell-cell fusion with wild-type *MAT***a** in a light dose-dependent manner and could reach comparable maximum levels as the wild-type *MAT*α emitter (**Fig. 2 C,D**).

Taken together, these results show that the opto-α strain is a suitable candidate, comparable to the wild-type, that produces physiological-range levels of pheromone production and mating behavior in response to light. Therefore, we employed the opto-α strain as a remotely controlled signal generator to create realistic α-factor spatial gradients within large populations of opto-α/MATa cells.

### 2.2 Optogenetic induction of pheromone degradation allows specific control of growth

It is expected that both pheromone production and pheromone degradation by the α-factor protease Bar1 shape extracellular pheromone gradients^30^. To gain spatial control over pheromone degradation, we constructed the opto-Bar1 *MAT***a** strain, in which the α-factor protease Bar1 expression is placed under control of the P_C120_ light-inducible promoter. We followed a similar promoter replacement strategy as for the opto-α strain, targeting the Bar1 locus in *MAT***a** cells (**Fig. 3A**). The expression levels and activity of Bar1 protein were measured using the *MAT***a** *bar1*Δ α-factor reporter, in the presence of purified α-factor added to the media. The levels of GFP produced by the reporter strain were used to quantify the production of Bar1, since Bar1 degrades purified α-factor in the culture medium (**Fig. 3A**). We observed light-dependent Bar1 activity, with the level of the reporter response decreasing as the light intensity increased (**Fig. 3B**). We reached higher Bar1 activity than the wild-type strain and undetectable basal (in the dark) Bar1 enzymatic activity.

**Figure 3.**
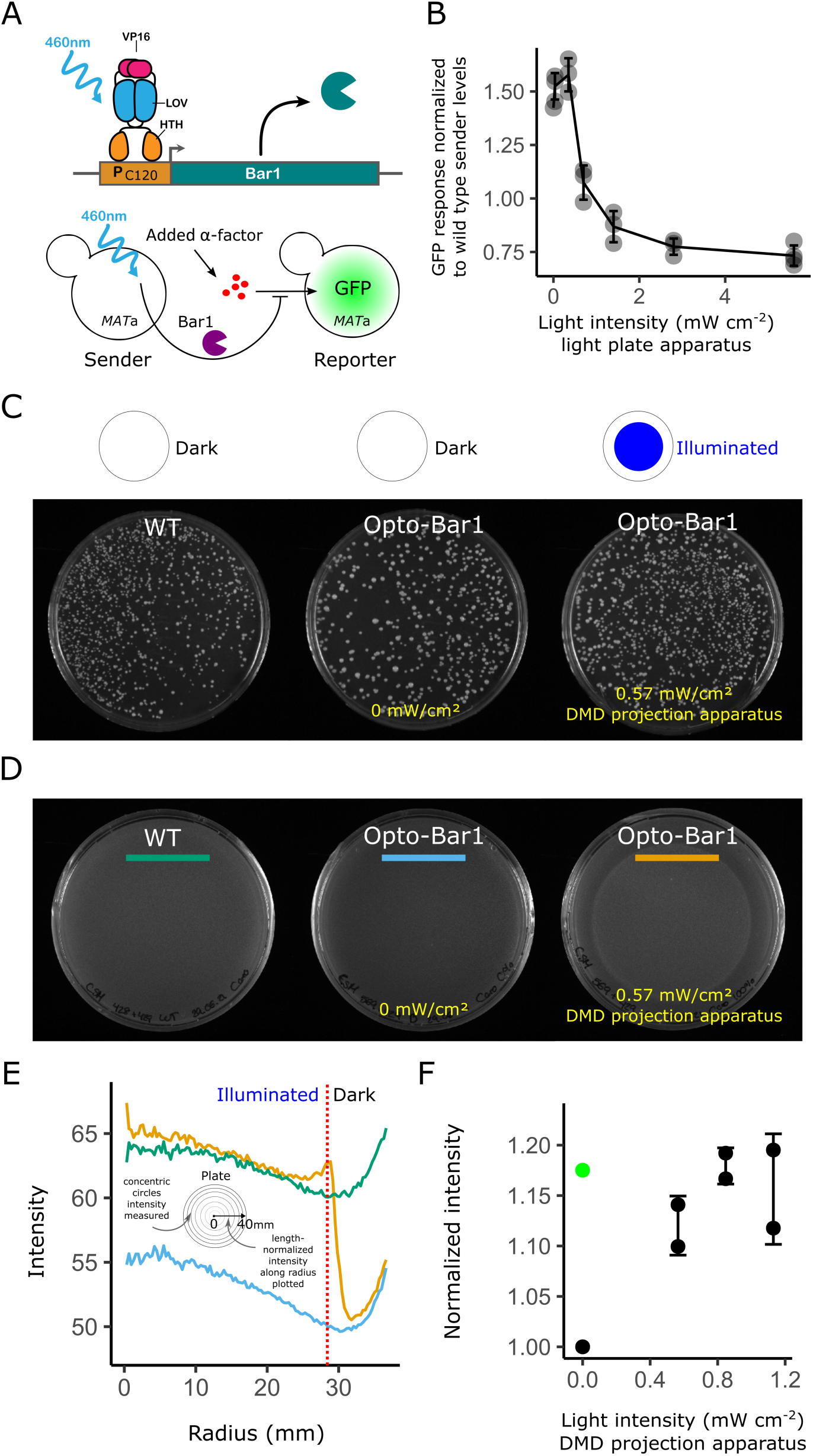
Optogenetic control of extracellular signal degradation allows tight spatial control of growth arrest. **A.** Promoter replacement in the Opto-Bar1 strain. EL222-mediated light-dependent expression of the native *loci* encoding Bar1 in the *MAT***a** strain (top) and the pheromone activity assay, in which exogenously added pheromone stimulates the sensitized GFP reporter strain (bottom). **B.** Resulting light-dose response from the assay, measured by flow cytometry. The biological activity of the pheromone is defined relative to the activity of the wild-type *MAT***a** strain. Error bars are the SD of 3 biological replicas**. C, D.** Diploid selective (C) and non-selective plates (D) showing mating efficiency and growth, respectively in illuminated and dark regions. **E**. Quantification of avoidance of cell cycle arrest (“intensity” corresponds pixel grey value which is proportional to yeast growth) in the Opto-Bar1 strain as a function of the distance from the center (a radius), based on the integrated pixel intensity of a radial profile (traces shown correspond to the colored bars in D, see methods). **F.** Light-dose dependency of cell cycle arrest in Petri-dish assays performed as in D. In this case, growth is normalized to the value of non-activated opto-Bar1 cells (lowest growth). Growth of the wild type is shown as a green dot. Error bars are the SD of two biological replicates.

Although Bar1 degrades the pheromone, its function has been long associated with increases in mating efficiency through several candidate mechanisms^14,26–29^ (see **Introduction**). However, the presence of Bar1 is not essential for mating itself. Indeed, when performing the optogenetic mating assay with the opto-Bar1 strain, cells located in dark regions—where *MAT***a** cells do not express Bar1—remain proficient at forming diploids (**Fig. 3C**), with a **∼**2-fold difference in mating efficiency between illuminated and dark regions (**Fig. S4**). Thus, contrary to optogenetic pheromone induction—and as expected—optogenetic pheromone degradation does not restrict but mildly enhances the final output of the pathway, namely the formation of diploids.

Cell cycle arrest, another physiological output of mating events, is controlled by the mating-pheromone pathway through the FAR1 branch^36^. When Bar1 is produced and actively degrades α-factor, one would expect to observe fewer cell cycle arrest events. Following similar methods as for diploid formation, we used the optogenetic plate assay to measure the capacity of *MAT***a** cells to escape cell cycle arrest as a function of Bar1 induction. We imaged an intermediate stage of the experiment (**Fig. 3D**). A differential growth pattern was observed in the non-selective plate used as template for replica plating in diploid-selective media (See also **Fig. S3**), as illuminated cells (expressing Bar1) grew similarly to wild-type levels, whereas non-illuminated cells did not grow significantly (**Fig. 3D-F).**

These results show that illuminated opto-bar1 cells recover the expected wild-type behavior in terms of α-factor degradation activity, mating, and proliferation. They also show that Bar1 is an effective spatial growth control actuator for synthetic biology applications.

### 2.3 Spatial optogenetic induction of α-factor production generates distance-dependent responses in mating populations

After quantitative characterization of our optogenetic strains, we next moved to assess the spatial properties of pheromone-mediated communication between *MAT***α** and *MAT***a** cells in an idealized (but physiological) context. To that end, we implemented the “half-domain assay”, which consists of monitoring *MAT*α/*MAT***a** mating reactions within a spatially homogeneous random distribution of cells of both mating types placed at the bottom of agarose-embedded “miniwells” (**Fig. 4A, see Methods**). Half of the total surface was exposed to blue light, to optogenetically induce pheromone production exclusively in the illuminated domain. The response of reporter cells along the axis perpendicular to the light-dark border allowed us to read and quantify the diffusion-reaction properties of α-factor.

**Figure 4.**
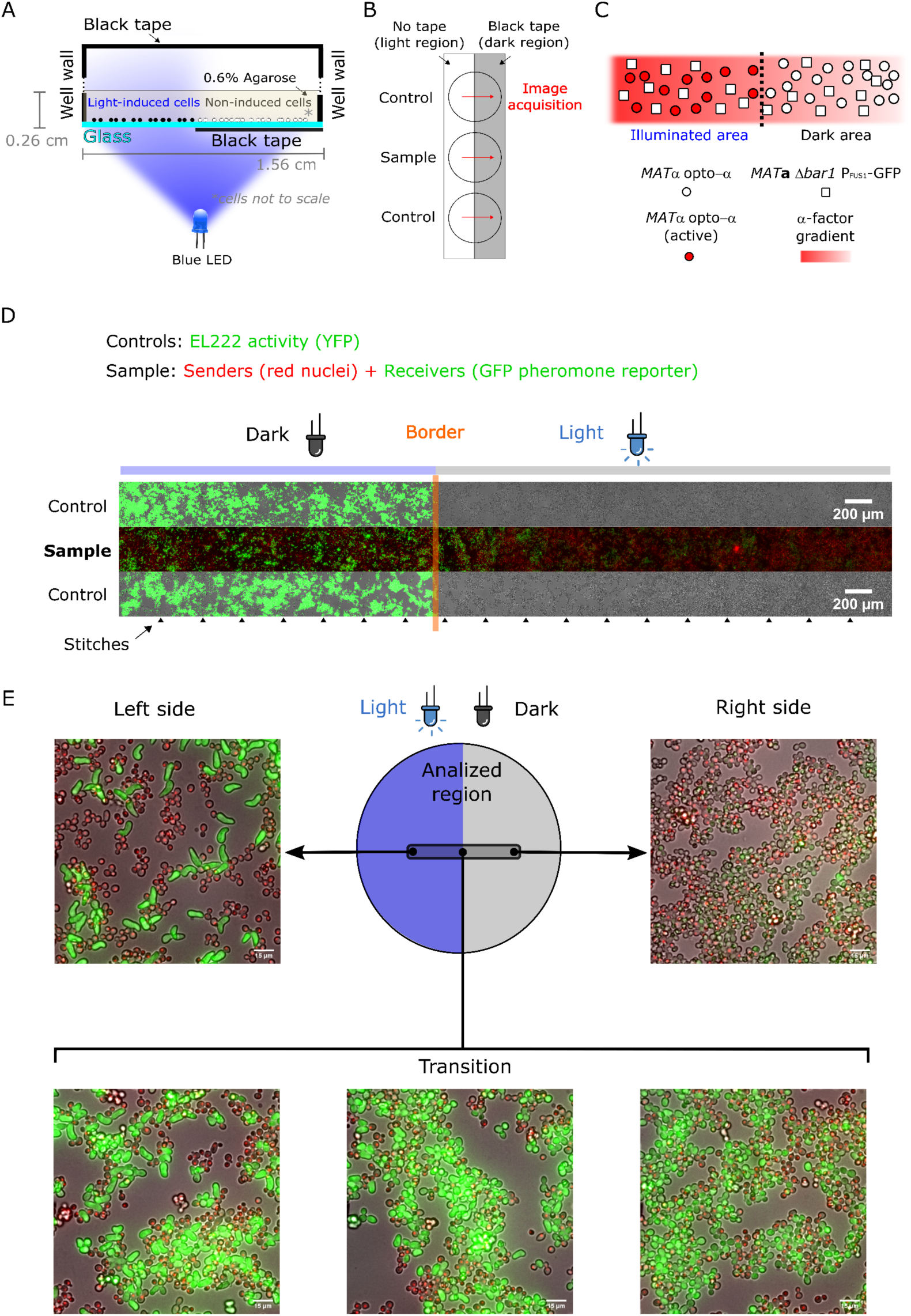
Optogenetic generation of pheromone gradients through the half-domain assay. **A.** Half-domain assay. Agarose-embedded cells are located at the bottom of the mini well and half of the glass surface is illuminated to induce EL222-dependent expression. **B.** To determine the position of the light border in samples along the acquisition path (red arrows), a control strain that activates EL222-dependent fluorescent protein expression is run in parallel in adjacent wells. The mean value of the border positions above and below the sample correspond to the position of the light border in the sample. **C.** Schematic representation of the experiment, close to the light-dark transition border. **D.** Microscopy image showing an overview of the mating reaction with the controls for the border location (top and bottom row) displaying strong YFP fluorescence on each side of the mating reaction (center row). Arrowheads at the bottom show where the image was stitched together. In the sample mating reaction (center), opto-α has a red nuclear marker while reporter *MAT***a** *bar1*Δ expresses GFP when stimulated by pheromone. Samples were imaged after 5 h of light exposure. **E.** Example fields along the experimental length showing the increased abundance of elongated phenotypes and the increase in GFP levels in the illuminated region.

We first conducted experiments with the opto-α strain as a light-dependent pheromone generator and the *bar1Δ MAT***a** reporter strain as a pheromone receiver and fluorescent reporter of mating behavior. Absence of Bar1 simulates natural situations where wild-type *MAT***a** receivers are in the minority (*e.g.*, upon a mating type switching event), where Bar1 levels are not enough to modify the global pheromone concentration. Thus, in this experiment, enzymatic degradation of α-factor by Bar1 is absent and disappearance of the pheromone is solely determined by its intake by cells^37^. After illumination (**See Methods**), cells along a stripe located in the center of the plate, with its long axis perpendicular to the light-dark interface, were imaged (**Fig. 4B)**. A schematic of the experiment and strains used is also shown (**Fig. 4C**). The opto-α strain was equipped with a red nuclear marker (the HTB2 histone fused to mApple) for easy identification, which is especially important in regions of clumping (**Fig. S5**, this new strain is named opto-α*). We used a 20x objective and an automated stitching method (**see Methods**) to obtain images of a large stripe (5671 µm x 330 µm) of both cell types using fluorescence and bright field microscopy at single-cell resolution (**Fig. S5**). Our experimental results revealed a sharp phenotypic transition between illuminated and non-illuminated areas. The sharpness of this transition (compared to the sharpness of the light border) was assessed by running a parallel experiment in neighboring mini-wells by stimulating a control strain that expresses YFP under the control of an equivalent EL222/P_C120_ light-inducible system. This control serves as a biological spatial imprint of the light-stimulated region, to both locate the position of the border in the sample and determine the sharpness of the light border (**Fig. 4D,E**). The region in the vicinity of the light-to-dark transition border illustrates the overall trend of the mating response in the *MAT***a** *bar1*Δ responder population: the GFP signal gradually vanished in cells further and further away from the frontier in the dark domain (**Fig. 4D**). Importantly, since we imaged cells at single-cell resolution, we clearly observed the cell morphology transitioned from elongated shapes to round as the distance from the light to dark frontier increased (**Fig. 4E**).

Our assay demonstrates that optogenetics can be used to produce large-scale population-generated gradients of a pheromone to infer the shape of the pheromone gradient through quantification of gene-expression and cell-morphology outputs (**Figs. 4C,D, S5).**

### 2.4 Biophysical modeling reveals a quantitative dependency of large-scale gradient shape on population-parameters

Using the half-domain assay, we next sought to infer the biophysical properties of the generated gradient. Thus, we constructed a biophysical model based on reaction-diffusion dynamics—as well as dose-response functions—and obtained single-cell quantifications of pheromone-responses using a combination of manual and automated single-cell segmentation and analysis (**see Methods**). We investigate what equation governs the time (*t*) evolution for the concentration of α-factor *c*(*x*, *z*, *t*) as a function of the position relative to the frontier of the dark domain (*x*, with the frontier at *x* = 0) and the distance from the glass surface (*z*, with the glass surface at *z* = 0). Pheromone-producing cells are located on the glass surface in our experiment, while pheromones diffuse into the agarose gel volume sitting on top of the cells (see **Fig. 5A**). Further, since we know that pheromone degradation happens mainly by cell uptake^37^ in a Bar1 knockout, degradation can happen only on the boundary close to the glass surface:

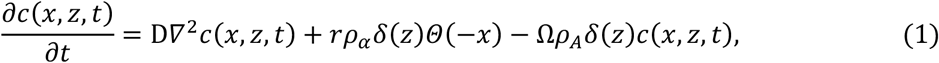

**Figure 5.**
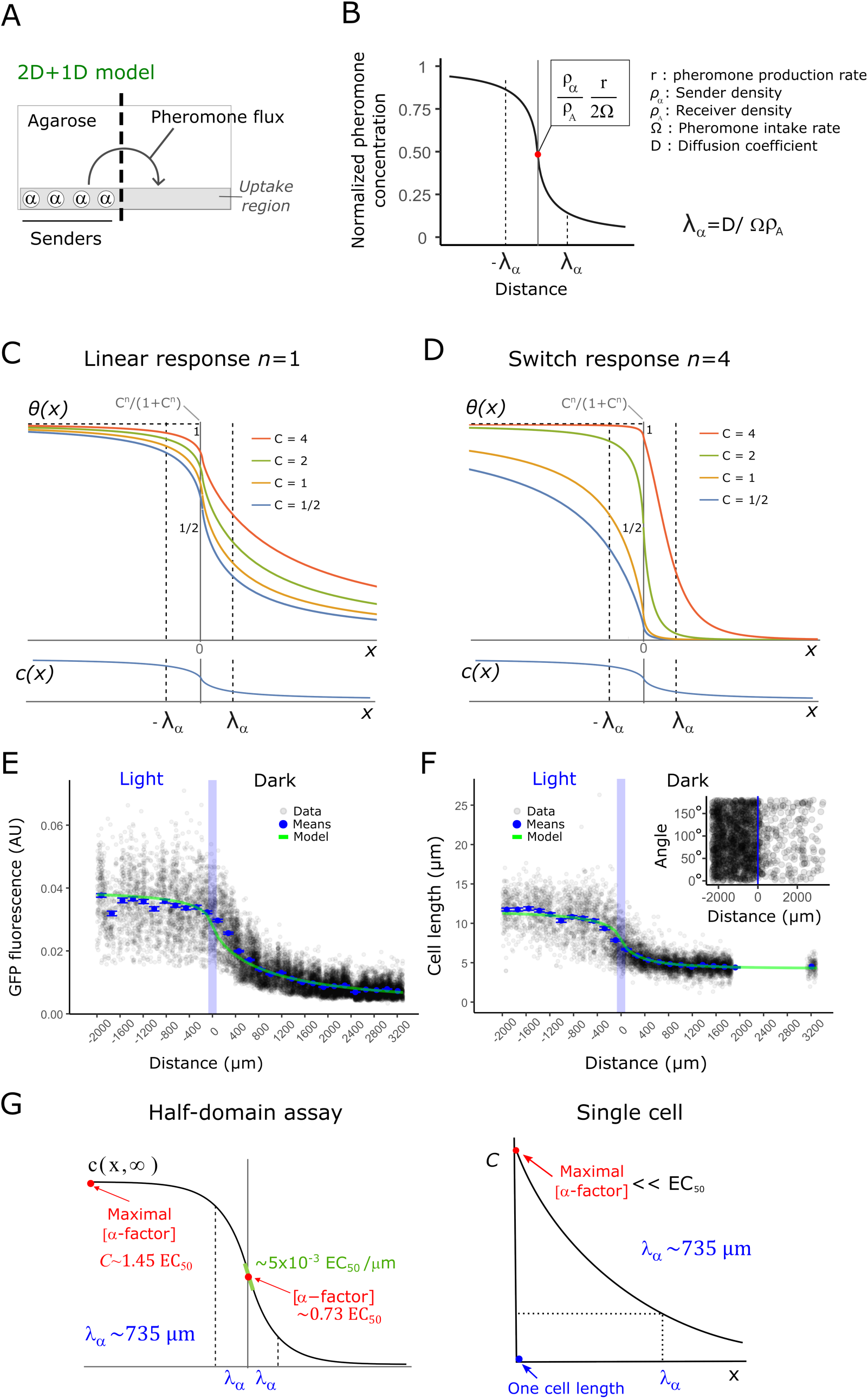
Quantitative properties of optogenetically generated gradients recapitulate switch-like and linear responses in *MAT*a cells. **A.** Schematic representation of the model used to infer α-factor gradient shape. The area on top of the cells is explicitly included and the region where pheromone uptake occurs is restricted to the bottom. **B.** Concentration profile of α-factor at steady state (Eq. 2, See **Supplementary Information**). **C, D.** Normalized response of the receptor (Eq. 3 normalized by its theoretical maximal value *C*^*n*^/(1 + *C*^*n*^)) for half-domain expression of α-factor, showing predictions for the shapes of responses using a Hill coefficient equal to *n*=1 (panel C) and *n*=4 (panel D). Hill coefficients greater than one correspond to a shift in the response towards a higher concentration of α-factor and a steeper response. **E.** Quantification of single-cell gene expression output (data: black, means: blue) and overlayed model fit (green), with the Hill coefficient set to 1, full set of parameters in Table S3. **F.** Quantification of single-cell gene morphological response (data: black, means: blue) and similarly overlayed model fit (green), with a fitted Hill coefficient of 3, full set of parameters in Table S3. The **inset** shows the absence of gradient alignment of *MAT*a *bar1*Δ. The orientation angle of single cells (points) with large area (a proxy for elongation in the light-domain) respect to the perpendicular to the light-dark transition border. An angle of 0° represents a cell with the long axis of its fitted ellipse oriented perpendicular to the light-dark transition border, while an angle of 90° the same axis is totally parallel. The cutoff for filtering out small cells was 200 pixels. **G.** Population-level and single-cell estimations of pheromone-profile parameters, based on the 2D+1 model.

which depends on its diffusion coefficient D, the cell surface density *ρ*_*α*_ and *ρ_A_* respectively for *MATα* and *MAT***a** cells, the pheromone production rate per cell *r*, and a parameter proportional to the rate of degradation of the pheromone per cell Ω, which has units of rate times length cubed. We analyze Ω further in the **Supplementary Information**. As an approximation, we assumed these parameters to be time-independent.

The solution to Eq. 1 at steady state is derived and plotted in **Supplementary Information,** the concentration is highest on the production domain far from the boundary and lowest on the diffusion domain symmetrically. The transition is logistic-like (**Fig. 5B**) and centered on the boundary, it is sharper closer to the glass surface and gradually smoother for larger values of *z* (see **Supplementary Information**). The precise shape of the transition depends on the finite size of the definition domain, specifically it depends on the length of the glass plate stripe *L_x_* and the thickness of the agar volume sitting on top of the cells **L*_z_*. Apart from these, the solution depends on a single length scale *λ*_*α*_ ≡ D/(Ω*ρ*_*A*_) defined by the rate between the diffusion and degradation rate density at the surface. In the large domain limit (*L_x_* ≫ *λ*_*α*_ and *L_z_* ≫ *λ*_*α*_) the finite size effects are negligible and the solution simplifies significantly. On the glass surface (*z* = 0) where the cells are located, the concentration is directly proportional to the ratio between production and degradation rates and can be described by following closed formula:

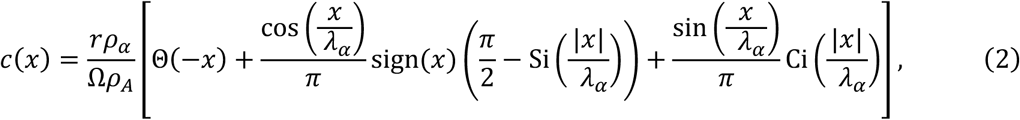

where Θ(*x*) and sign(*x*) are respectively the step function and the sign of its argument while Ci(*x*) and Si(*x*) are the trigonometric integrals functions. While Eq. 2 is logistic-like in shape, it features a heavy tailed decay of the gradients (as *c*(*x*) ∝ *λ*_*α*_/*x*) for large distances from the light boundary (*x* ≫ *λ*_*α*_), in sharp contrast to systems traditionally modelled in one-dimension—such as the drosophila embryo—where decays are exponential. The concentration at the boundary *c*(0) = *rρ*_*α*_/(2Ω*ρ*_*A*_) depends on the rates of emission and absorption as well as the surface density of emitter and absorbing cells. On the glass surface in this half domain essay, we expect densities of α-factor much greater than the one obtainable for a single cell-source, since the density of producers is greater than one cell within one decay length of the observed profiles (*ρ*_*α*_ > 1/*λ*_*α*_^2^).

To connect these analytical results to the experiments, the response *θ*(*x*) of the mating pheromone pathway to the steady-state concentration of pheromone is modeled by composing the result for *c*(*x*) with a Hill function with coefficient *n*. This results in:

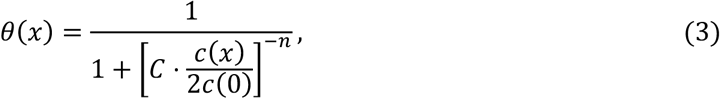

where *C* ≡ 2*c*(0)/EC_50_ is the concentration of the ligand far away from the border (*x* → −∞) in “EC_50_” units, namely, the concentration producing the half-maximal P*_FUS1_* mediated gene-expression response. We obtained the theoretical shape for a standard concentration profile at various values of *C* and two different values of *n* **(Fig. 5C,D)**. It is well known that no cooperativity is involved in the gene expression output from the P_FUS1_ promoter^14^, we thus fixed *n* = 1 for gene expression output. Experimental responses are measured in unknown concentration units and are affected by a baseline that needs to be determined, for this reason we fit the experimental data to a formula that takes into account these two extra parameters 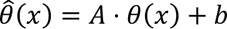. It is important to note that the Hill-equations saturate at very large concentrations (*C* ≫ EC_50_), while in all our experiments we determined *C* (the pheromone concentration at the curve’s plateau) to be of the same order of magnitude as its EC_50_, *i.e.* the response curve becomes flat far from the border in the left domain because of a constant concentration of pheromone, and not because of saturation (see Section 2.5 below). We were able to fit independently both the normalization parameter *A* as well as the ligand concentration far away from the border *C* thanks to the availability of two readouts of the pheromone concentration with different Hill coefficients *n*.

### 2.5 Quantitative analysis reveals that population-level pheromone gradients are long-range, shallow, and limit morphological transitions to the vicinity of the source

Armed with the model derived in the previous section, we quantified the single-cell outputs of the activity of the mating pheromone pathway—specifically, P*_FUS1_* dependent gene-expression (**Fig. 5E**; determined using standard illumination correction and segmentation tools; **see Methods**) and cell elongation (**Fig. 5F**; determined independently and manually; **see Methods**)—in *MAT***a** cells lacking Bar1. Our results revealed a gradient of fluorescence intensity that extends far beyond the region where the pheromone is generated (**Fig. 5E**). Thus, the transport properties of α-factor can generate responses at distances spanning hundreds of cell lengths, consistent with a widely diffusible molecule. The morphological transition (**Fig. 5F**) is steeper than the gene expression response, with cells increasing their maximal length within a narrower distance range and at a higher pheromone concentration. This is expected, as the morphological transition is indeed switch-like^18,20^.

We ran control experiments using (i) opto-α emitters mixed with the wild-type *MAT***a** reporter (expressing Bar1) and (ii) a mixture of the wild-type *MAT*α as light-independent emitters and the *MAT***a** *bar1Δ* reporter. To ease cell segmentation, we performed a machine-learning based protocol (See methods) to obtain a global picture of the behavior of both samples and controls. In the reaction with wild-type expression of Bar1, basal GFP output was obtained with no detectable differences between the light and the dark domains. In contrast, in the mix with wild-type *MAT*α and the *MAT***a** *bar1Δ* reporter, strong activation was observed in both illuminated and dark domains (**Fig. S6**). These controls further allowed us to compare the maximum GFP output activation of the opto-α and wild-type *MAT*α strains to confirm that the emission levels are close to physiological in the half-domain assay, as well as allowing us to correct illumination inhomogeneities and background-bleaching effects of light induction with an independent method (**Fig. S6, see Methods**).

We also observed that cells elongate, but do not preferentially grow towards (nor away from) the large-scale gradient, as revealed by the lack of enrichment of cells with orientations close to 0° or 180° respect to the direction of the gradient (**Fig. 5F, inset**), demonstrating that—even at its steepest point (the border)—the large-scale gradient is insufficient to induce chemotropism. In addition, because we did not observe formation of single or multiple “thin protrusions”, characteristic of saturation of the pheromone pathway and observed in our reporter^14,19,38^, we concluded that cell responses lie within the sensitive range of the pathway, which is indicative of the Ste2 receptor not being saturated.

Next, we used the biophysical model (**see Supplementary information**) to infer key quantitative parameters of the gradient. We fitted the experimental datapoints for the two observables to 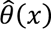 (**Fig. S7**), with *n* fixed to one for the gene expression output (**see Methods**), and fits displayed on top of the raw rather than the normalized data (**Fig. 5E,F**). In this way, we inferred the basic properties of the gradients generated (**Fig. 5G**): the best fit returned us the steady-state decay length *λ*_*α*_∼ 735 µm, the steady state concentration of α-factor for *x* ≪ −*λ*_*α*_, *C* = 1.45 EC_50_, and the Hill coefficient of the morphological response (*n* = 3). The gradient value at light-dark transition border, the expected steepest point of the large-scale gradient, has a value of ∼ 5 ⋅ 10^&*^ EC_50_ /µm. Using an EC_50_ value for the response of 1‒3 nM^14,18,19^, we obtain a value of 5 to 25 pM/µm or, roughly, ∼70 pM per cell diameter (one cell diameter is ∼5µm). Such shallow gradients will produce differences in receptor occupancy between each side of the cell below the minimal ∼0.5% observed in chemotropic cells in artificial pheromone gradients^39^.

Collectively, these results imply—in this spatial arrangement and in the absence of Bar1—that: *(i)* pheromone levels characteristically decay roughly ∼100 cell bodies away from producer regions or single cells; *(ii)* maximal steady-state pheromone levels found in typical (dense but not crowded) mating populations are not saturating and rather very similar (less than double) to the EC_50_of the response (its most sensitive point); *(iii)* cells “ignore” (filter out) large-scale gradients (such as those produced within the spatial limit of finite emitter populations) that are non-informative about the location of nearby emitters; and *(iv)* knowing the dependency of pheromone concentration at the border on the linear density of cells (∼100 cells/mm or ∼70 cells in one decay length), *C*/2 would be reduced by around two orders of magnitude for a single cell located at the border. This means that for a single cell, pheromone levels close to the EC_50_ cannot be reached, even in its immediate vicinity. This is consistent with the need for close-proximity mechanisms and focal pheromone emission^9^ for such close-contact situations. In other words, single emitters cannot induce elongation or chemotropism in responding cells, even in the absence of Bar1. Sufficient pheromone levels to induce mating behavior at a distance are always produced by a collective of cells and under low Bar1 conditions (biased ratios). Increased rates of degradation of the pheromone (*e.g.*, in the presence of *MAT***a**-bound Bar1) would scale down the gradient. These constraints over the maximum levels of pheromone that a single cell can produce might explain why—when homogeneous Bar1 sources are present—we did not observe elongation in either of the two domains (**Fig. S6**) and the response values stayed at basal levels, despite the high density of emitters.

### 2.6 Quantitative analysis of large-scale gradients produced by opto-Bar1 receivers suggests localization of Bar1 activity in its production domain

Our experiments in the absence of Bar1 are adequate to explain natural situations where wild-type *MAT***a** receivers are in the minority (*e.g.*, upon a mating type switching event), such that Bar1 is not abundant enough to modify the large-scale pheromone landscape. As the mating-type ratio becomes unbiased, *MAT***a**-produced Bar1 protease becomes important. When mated under conditions of pure diffusion-reaction—but not advection—Bar1 is believed to allow reshaping of the pheromone landscape. The exact nature and biological function of this reshaping activity during mating remains controversial^3^. Therefore, we used the half-domain assay with the opto-Bar1 strain in the presence of wild-type MATα cells as emitters to shed light on the problem.

In this case, WT pheromone production is homogeneously distributed on the glass surface and emission of Bar1 is localized in one half-domain of the glass surface (**Fig. 6A**), on the right of the light-dark transition border (**Fig. 6A, Fig. S8**). For this assay, wild-type *MAT*α emitter cells were tagged with a green nuclear tag, while opto-Bar1 cells were tagged with a red nuclear tag. This allowed us to confirm that, as in the Petri dish experiments (**Fig. 3D-F**, **Fig. S4**), mating events (measured as nuclei fusion events) happen in both illuminated and non-illuminated areas and that Bar1-mediated increases in mating events correlate with the increased initial growth of *MAT***a** (**Fig. S9**). Crucially, observations of the cell-elongation phenotype revealed a striking difference compared to this response’s profile in the opto-α experiment (**Fig. 5)**: in the opto-Bar1 case, the elongation pattern emerges far away from the border, well into the dark (left) region of the experiment (**Fig. 6B-C**).

**Figure 6.**
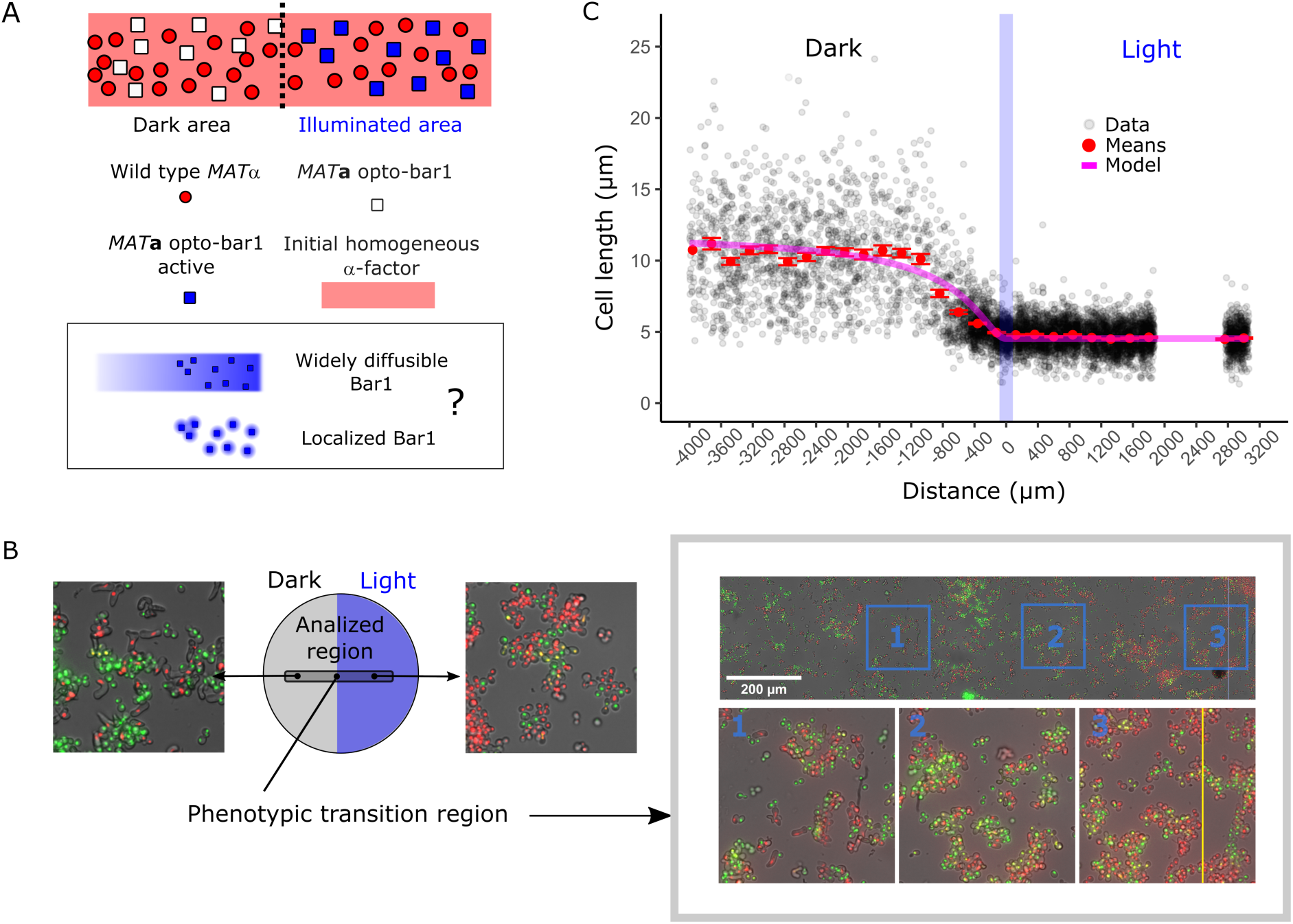
Bar1-generated α-factor gradients reveals restricted Bar1 diffusion. **A.** Schematic representation of the half-domain assay for opto-Bar1, close to the light-dark transition border. Emitter cells produce α-factor homogeneously. Light-induced Bar1 secretion occurs with unknown spatial distributions (bottom box). **B.** Zoomed-out depiction of the half-domain assay for Bar1 production and microscopy imaging of Bar1 emitters (cells with red nuclei), receivers (cells with green nuclei), and newly formed diploids (cells with yellow nuclei). The yellow line represents the position of the left limit of the light-dark transition border. **C.** Quantification of the single-cell morphological response (cell length) of opto-Bar1. The light border is represented by the blue vertical line corresponding to the border location determined with controls run in parallel (See Methods). The blue dots correspond to the mean value in each one of 30 bins and the error bar represents its standard error. The purple line represents the zero parameter fit of the perfect sink model (see text).

To understand the observation we turned to modeling, and—as a simplification—we solved the Bar1 production domain acting as a perfect sink for α-factor model. In this configuration, Bar1 is confined to the glass surface in the production domain (*z* = 0, *x* > 0) and the *α*-factor degradation rate by Bar1 is set so high that it effectively degrades all *α*-factor in its presence. These limits lend themselves to a simple change in boundary conditions at *z* = 0, for the diffusion of α-factor, respect to Eq. 1 (derivations and detailed solution **Supplementary Information**). Similarly to what has been used to obtain Eq. 2, in the infinite domain limit the profile for α-factor concentration on the glass domain follows the following:

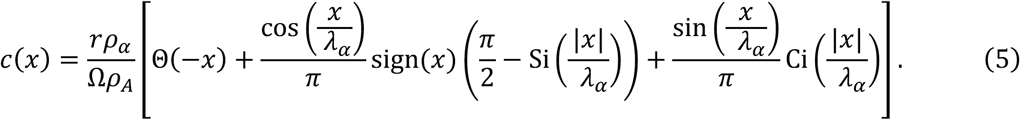

Even if α-factor is produced on the full glass plate (*vs.* Eq. 2 where α-factor is produced only on a half domain), due to the high efficiency of Bar1 in degrading α-factor (*vs.* Eq. 2 where α-factor is degraded by cell intake), in Eq. 5, *c*(*x*) is exactly equal to zero over the full Bar1 production domain. Further, due to the confinement of Bar1 on the glass surface of the producing domain, *c*(*x*) quickly increases on the diffusion domain with a slope that is set by the same *λ*_*α*_ that has been measured in the experiment without any Bar1. Finally, for the same reason, the concentration far from the boundary 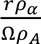 as well as the Hill coefficient *n*, the baseline *B* and the arbitrary unit *A*, in the α-factor concentration dependent cell response function 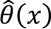 are predicted to be the same as in the experiment without any Bar1. Considering this, we overlapped the predicted cell response function for elongation 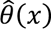, using the solution to Eq. 5 for the α-factor concentration, to the experimental data in presence of Bar1, fixing all the parameters using the ones fitted to the experiments in absence of Bar1 presented in the previous section (*i.e.* with zero free parameters) and obtained a reasonable agreement to experimental data (**Fig. 6C**). This indicates that Bar1 is efficient in degrading α-factor respect to cell intake, and that its diffusion is compatible with localization on the glass domain in the production domain. Further, if we allowed in the equations for Bar1 to diffuse substantially in the agar volume above the producing cell, due to the long range nature of diffusion in our geometry, and the progressive flattening up of the gradients above the glass surface (see Eq. 2), Bar1 would have penetrated substantially in the diffusion domain and it would have degraded α-factor there substantially reducing the cellular response respect to the experiment without Bar1. Considering that this didn’t happen, we conclude that Bar1 penetration is limited.

## 3. Discussion

Using our optogenetic tool, here we show that in sparse yeast populations, biologically relevant concentrations of α-factor in the environment are built up by the contributions of many emitter cells, and that there is a length-scale for gradient formation other than that in the well-known immediate vicinity of cells (1 cell body or ∼5 µm); namely, the large-scale gradient formed at the edge of finite producer populations. Chemotropism directed along these collectively generated large-scale gradients was not observed in our half-domain assays. These large-scale gradients are thus not steep enough to induce chemotropism, despite pheromone concentrations being high enough to trigger strong elongation. Then, *MAT***a** cells filter out these uninformative large-scale gradients and rather only follow local gradients with sufficient steepness^9^. Elongation responses in this case better resembles random rather than active search for mates.

Our model confirms previous calculations^8^ showing that concentrations of pheromone emanating from a single emitter must be well below the mating threshold (**Fig 5H**), this means that pheromone secretion by a single cell doesn’t generate meaningful long-range gradients that can be followed with chemotropism involving cell elongation—even if partners are sensitized (without expressing Bar1). Long-range gradients are therefore only collectively produced in mating reactions. Indeed, compared to values generating long-range chemotropism in microfluidic devices, large scale gradients in the half-domain assays are roughly ∼10x shallower (we report ∼5 to 25 pM/µm whereas in studies with precise quantification of the gradients generated, these are in the order of hundredths of pM/µm^19,40^). We expect very similar behavior for wild type α-factor emitter, because the pheromone production rates are comparatively very similar.

Our results also show that Bar1 effectively tends to stay in its production domain. It is possible that such behavior comes from out-of-steady-state effects. Namely, that both pheromone and Bar1 diffuse widely, but that Bar1, due to its size, does so slower. Taking the values for diffusion from earlier simulation work^30^ we get half a millimeter diffusion on the timescale of early cell responses (45 minutes), which is ballpark compatible with our observations of a shift of the boundary into the left domain. On the other hand, it is well known that Bar1 is an abundant disulfide-bridged mannoprotein present in the cell wall of *MAT***a** cells^41^ and this sink character is essential to build models that explain the establishment of gradient disentanglement from two sources in close-proximity intra-ascus interactions^28^ and mutual-avoidance in *MAT***a** cells^29^. Simulation work has either introduce high bulk degradation of Bar1^29^ or zero-diffusion of Bar1 to keep Bar1 halos localized to emitter cells. Considering the clear evidence for Bar1 association with cell walls and the lack of evidence for specific Bar1 degradation, the latter alternative—added to slower diffusion for the freely diffusible Bar1—seems the most likely explanation. More experimentation is needed to elucidate the precise mobility of Bar1 during mating.

In any case, such limited diffusion of Bar1 does not prevent *MAT***a** from modifying global pheromone profiles—provided cells are at a sufficient cell density. Indeed, in our opto-Bar1 experiment (**Fig. 6A-C**), Bar1 greatly reduces the global α-factor concentration, and *MAT***a** cells do not elongate nor arrest (**Fig S6**). Despite this, cells in close contact still mate normally (**Fig. S9**). Thus, when separated in space, cells cannot reach the threshold for executing the morphological transition. This is in agreement with the absence of elongation and the persistence of cell division by *MAT***a** cells in unbiased mating reactions in the presence of advection^15^. In this sub-threshold case, it is likely that increased cell-number, randomly oriented bipolar budding^10,15^ and close-contact mechanisms^9,42^ aid mating. These known mechanisms are functionally very different as search strategies to what we here call long-range chemotropism, a biological function first inferred from the classical experiments by Segall in the early 1990s^24^, in which cell-cycle arrested cells in agarose extend their cell bodies towards a glass pipet that releases pheromone. This type of chemotropism, involving cell elongation, is only accurate at concentrations matching those at which cells switch^43–46^. Therefore, gradients need to fulfill two conditions to induce this behavior: they must have a high mean pheromone concentration to induce elongation and need to be steep enough to direct the polarisome towards the source. Our results suggest that—in mating reactions with cell densities that would render chemotropic cell elongation useful—there is a trade-off between the mean pheromone concentration and the steepness of the gradient: cells are weak producers but, in principle, good position signalers, whereas dense populations are strong signalers but weak position signalers.

Thus, our results explain why elongated *MAT***a** chemotropism is rarely observed as directed mate-searching in mating reactions with sparse populations. Indeed, depending on the levels of Bar1, within environments crowded with α-factor emitters, cells will either restrict pheromone levels to the emitter vicinity and rely on local (short-range chemotropism) mechanisms^9,42^ or exhibit randomly oriented elongation due to gradient overlapping. Randomly oriented cell elongation could be a useful strategy under such low Bar1 conditions, as gradient overlapping will make the switch concentration proportional to the cell density of partners, rather than their distances. Our results reveal that gradient overlapping is significant in the absence of Bar1, implying that distance-to-source measurements by cells in the production region cannot be disentangled from demographic (population-parameter) measurements. Therefore, our results highlight a limitation of long-range chemotropism as an efficient mate-directed search strategy; namely, the necessity of the *MAT*α-biased ratios for *MAT***a** to reach its switching threshold entails an increased likelihood of producing local, uninformative gradients caused by increased overlapping. This trade-off, in principle, could be balanced at some specific ratio, where Bar1 levels are enough to spatially restrict levels of α-factor and sources are clustered to allow a high local concentration of α-factor. This could be the case, for example, for a larger patch of *MAT*α encountering a close smaller patch of *MAT***a** after mating-type switching^4^, spore dispersal in wild isolates with delayed mating interest^12,47^, or ordered seeding for patterning applications^48,49^.

Overall, our results suggest that the pheromone landscapes in sparse mating reactions are directly shaped by population parameters and differential diffusion of Bar1 and α-factor observations. Functionally, steep gradients localized at either the source or the sink can only guide chemotropism when the population ratios are biased enough to limit global Bar1 availability. As suggested by others, due to its sink function, the gradients might be steeper at a receiver cell; thus, if several receiver cells cluster, the gradients can direct these cells away from each other^29^. Thus, gradients can effectively provide information about the location of competitors (other *MAT***a** cells) rather than potential mates (*MAT*α cells). In addition, local degradation of α-factor allows the asexual proliferation of unmated individuals, maximizing the total population’s fitness.

We expect our optogenetic tools to be useful to prove the limits of successful chemotropism—where cells fuse—in microfluidic confinement of small groups of cells and/or microcolonies stimulated with high spatial illumination resolution. In the same context, controlling dynamically (taking advantage of EL222’s reversibility^50^) the levels of Bar1 and pheromones (eventually including a-factor optogenetic control), in confinement, could allow a deeper quantitative understanding and controllability of mate search (and self-avoidance^51^). Controlling cell-cell fusion precisely in space could be also useful in the context of engineered living materials^52^

Our study encourages the use of the yeast mating system as a quantitative framework for the design and control of spatial behavior of natural and synthetic cell-cell communication systems.

## 4. Methods

All materials described here and in the Supplementary Information are available upon reasonable request. Datasets used to generate the figures and the code for the mathematical modeling can be found on the public Zenodo archive 10.5281/zenodo.10572403

### 4.1 Strains and growth conditions

*S. cerevisiae* strains used in this study (**Supplementary Table 1**) are derivatives of BY4741 (*MAT***a** *his3*Δ*1 leu2*Δ*0 met15*Δ*0*), BY4742 (MATα, *lys2*Δ*0,* otherwise identical to BY4741), SEY6210a^53^ (*MAT***a** *leu2-3,112 ura3-52 his3*Δ*200 trp1*Δ*901 lys2-801 suc2Δ9*), and SEY6210 (MATα, otherwise identical to SEY6210a). Yeast cells were routinely grown on complete medium (Yeast peptone dextrose; YPD, Sigma) or synthetic complete (SC) medium—composed of 6.7 g Yeast Nitrogen Base without amino acids (Difco 291940) and 0.8 g complete supplement mixture drop-out (Formedium DCS0019) in 1 L—supplemented with 2% glucose (Merck). Cells from glycerol stocks or agar plates were inoculated into YPD and incubated at 30°C on an orbital shaker at 250 rpm (in an Innova 4230 incubator) for 12 to 16 h. Day cultures were performed by inoculating 5 mL of SC or YPD media with overnight cultures at a concentration of 1% *v/v*. Care was taken to reduce unwanted light exposure as much as possible before the start of the experiments by covering the agar plates and precultures in aluminum foil and by performing all steps without direct exposure to light (low ambient light).

All yeast transformations were conducted following a standard lithium acetate protocol. To replace the native promoters located upstream of the genes of interest with the EL222-dependent promoter P_C120_, we performed CRISPR/Cas9-based genome editing following a method described previously^54^. Oligonucleotides used in this study are listed in **Supplementary Table 2**. The replacement fragments used to exchange the upstream region of the relevant genes (pheromones or Bar1) were amplified from plasmid pYTK097, which contains the original sequence of the wild-type P_C120_ promoter from *Erythrobacter litoralis*^55^, using primers containing appropriate homology regions as overhangs. DNA sequences coding for guide-RNA (gRNA) sequences (obtained from IDT) used to target Cas9 to the desired loci are also listed in **Supplementary Table 2**. Short and long gRNA were hybridized with each other and ligated into plasmid pML104, which possesses a *Cas9* expression cassette and an URA3 marker for auxotrophic selection. The EL222 transcription factor was inserted at the HIS3 locus by digesting plasmid pEZ-L105^56^ with the *SfiI* restriction enzyme. In this construct, EL222 is expressed from the constitutive P_pgk1_ promoter. Nuclear tagging was performed by inserting mApple at the HTB2 locus by amplifying mApple from plasmid pPH330 using primers oPH_0386 and oPH_0387 followed by transformation and selection using G418 antibiotic selection. Nuclear tagging with mVenus was performed by replacing mApple in the strain above using CRISPR/Cas9. The mVenus fragment was amplified from plasmid pPH_086 using the same oPH_0944 and oPH_0945 primers and guideRNAs oPH_941 and oPH_942.

### 4.3 Light stimulation experiments in 24-well-plates

Overnight cultures were diluted 1:50 in fresh LoFlo-SC (synthetic complete media prepared with “low fluorescence” yeast nitrogen base lacking folic acid and riboflavin; Formedium) to reduce media fluorescence and grown to an optical density (OD_600_) of 0.4‒0.5. Individual day cultures of optogenetic and reporter strains were washed once with 1 volume of LoFlo-SC media, resuspended to an OD of 0.5, mixed at a 1:1 ratio, and placed in black glass-bottom 24-well plates (CellVis). Light stimulation was performed in a custom-made light plate apparatus based on the design by Gerhardt et al^33^ equipped with a blue LED (λ=461 nm). The final volume in all experiments was 1 mL and the incubation time was 2 h. When opto-Bar1 was used as a test strain, α-factor (Sigma) was added at a final concentration of 1 µM. A well containing only the reporter strain was routinely included to calculate the basal GFP expression from the P_C120_ promoter. Light intensity calibration of the apparatus was performed as reported before^33^ using a power-meter and a custom 3D-printed adaptor. Reporter gene expression was analyzed by flow cytometry immediately after sampling.

### 4.4 Patterned light stimulation in Petri plates

Day cultures in SD media containing all amino acids were inoculated at 4% *v/v* with an overnight culture and grown until the OD_600_ was ∼1.0The cultures were then centrifuged at 3500 rpm for 7 min, the supernatant removed, the pellets washed once in SD medium containing all amino acids, and the optical density adjusted to OD_600_ =1.0. These suspensions were mixed at a 1:1 ratio and 150 µL was added to 6 mL of 0.5% agar (“top” agar, kept in a water bath at 55°C).

For the “velvet” version of the assay (**Figs. 3, S4, S5**), the top agar was poured over Petri plates containing solid SC-agar (1.5% agar; 20 mL total volume) supplemented with all amino acids., using a leveling surface. Plates were then sealed with parafilm, keeping the lids of the plates slightly tilted. Blue light patterning was performed with a DMD projector (DLP® LightCrafter™ 4500 TexasInstrument) placed inside an incubator (Memmert UF160); the projector projects light of the desired shape and intensity over Petri plates placed over a black surface. After light stimulation, the plates were replicated on SD plates lacking methionine and lysine using the sterile velvet technique, incubated for at least 24 h at 30°C, and imaged using a multi-imager (Biorad) for colony counting.

For the “direct” version of the assay (**Figs. 2, S4, S6**), agar cell suspension was poured over 20 mL of 1.5% agar SD media with low lysin levels (0.005 g/L) and was directly recovered for imaging and colony counting after at least 24 h of incubation (**Fig. S4**). The basis for this simpler version of the assay relies on the following: because the *MAT*a and *MAT*α wild-type strains are auxotrophic for methionine and lysine, respectively (**See Supplementary Table 1**), we expected haploid growth— and presumably mating—to depend on supplementation of these amino acids. Consequently, a low concentration of these amino acids would allow mating, but later only select for diploid growth as these amino acids become exhausted. We note that the methionine levels in the 150 µL cell mixes are enough to allow mating, as adding methionine did not increase diploid growth. On the contrary, lysine levels were optimized to determine the final concentration used (levels higher than 0.005 g/L did not affect diploid growth). Calibration of the light intensity of the device was performed by placing a power meter at the location of the agar surface. For the determination of growth levels, we used the *radial profile plot* function of Fiji, where the intensity at any given distance from the central point represents the sum of the pixel values around a circle (**Fig. 3F**, inset). Such integrated intensity is divided by the number of pixels in the circle, yielding normalized comparable values.

### 4.5 Half-domain gradient generation assay

Rows of wells in glass-bottom 24-well plates (CellVis) were taped in the “row” direction following available marks in the frame that coincide with the hemisphere of the wells. Longitudinal sections of black tape were cut and the uncut side of the tape was gently positioned facing the hemisphere of the wells. Overnight cultures were reinoculated at 2% *v/v* in YPD. These day cultures were grown for 4 h, washed once in 4 mL of synthetic complete media with 2% glc (SC-GLC), and then resuspended in 1 mL of the same media. The OD600 was determined in a cuvette benchtop spectrophotometer using 100 µL of this culture and 900 µL of SD-glc. Cultures were diluted back to OD_600_=0.1 in a final volume of 5 mL **SC-glc**. Concanavalin A coating was performed as follows: wells were exposed to 300 µL of 0.2 mg/mL Concanavalin A (ConA, Sigma; 1/10 dilution of a 2 mg/mL stock that was spun down to remove protein aggregates) for 6 minutes, then the solution was removed from each well (using a the P1000 pipet and then a P200 for the remanent), washed once with 300 µL of sterile water (water was removed in one strong inversion movement and not pipetted out), then 300 µL of the appropriate cell suspension mixes were gently deposited (consistently on the west side of each well) and allowed to settle gravitationally for 35 min. During cell settlement, fresh 1.5% *w/v* low-melting point agarose (Sigma) was melted in SD-glc in a glass beaker with constant and gentle magnetic stirring. Then, 200 µL of the hot (70°C) agarose solution was taken out (reverse pipetted) with a 1000 µL (blue) tip, held for 10 seconds, and the solution was gently (5 seconds to dispense 200 µL) was deposited in the bottom “west” position, immediately followed by gently pipetting up and down (close to the middle high, also pointing in the west direction). We set a light exposure time of 5 h, which allowed us to observe complete development of the output curves. The total analyzed region was roughly 5.4 mm around the border. Wells have a diameter of 1.554 cm, and cells are embedded in a total volume of 0.5 cm^3^ of agarose.

### 4.6 Flow cytometry

Flow cytometry measurements were performed on an LSR II cytometer (Becton-Dickinson). GFP fluorescence was measured with a 488 nm laser using BD FACS DIVA software (BD Biosciences). First, cells were gated in an FSC-A/SSC-to exclude debris and doublets. GFP-positive cells (either *MAT***a** or *MAT*α, depending on the experiment) were gated manually using FlowJo (Becton-Dickinson), and the mean fluorescence of this population was used for further analysis.

### 4.7 Fluorescence microscopy

Imaging of the half-domain assay was performed using a Leica DMi8 inverted microscope (Leica Microsystems) equipped with a motorized stage and a 20x objective (N.A. 0.4) using *LasX* software (Leica Microsystems) in multi-position mode. Fluorescence images were acquired using the 470/40 (Ex), 525/50 (EM) filter for GFP and the TXR filter 560/40 (EX), 630/75 (EM) filter for mApple proteins, respectively. Imaging was also performed using an inverted Olympus IX81 epifluorescence microscope equipped with an *“*Xcite exacte*”* light source (I = 30%) and a filter cube (EX) 514 nm/10 (EM) 545 nm/40 (49905 – ET) with a 1000 ms exposure time.

### 4.8 Manual determination of cell-length

As a proxy for the morphological response to pheromone we manually measured the cell-length for cells in the half-domain experiment. In the opto-α experiment, a total of 6995 cells were segmented, while in the oplto-Bar1 experiment the total was 8331 cells (a larger region was sampled in this experiment). In the range plotted the total numbers analyzed correspond to 6686 and 7231 cells, respectively. A small fraction of the cells at the leftmost border is taken out to avoid reflective border effects on the accumulation of α-factor (which are not modelled). Lengths were measured manually using the segmented line tool in Fiji in single cells. On each identified cell, we measured its length by drawing a line along its longest diameter. In elongated cells we draw their central longitudinal axis using as many points as needed to maximize precision (elongated cells are curved). Although we counted mostly non-overlapping cells, if a mild overlap existed, or the overlap produced clearly distinguishable boundaries, cells were considered. Prezygotes (cells having a characteristic dumbbell shape, product of cell-cell fusion events), were not considered in the analysis. In samples with large cell-clumps, we measured regions close to empty spaces where little overlap existed. All cell segmentation data and original images are available in Zenodo.

### 4.9 Machine-learning based cell segmentation

Manual segmentation is very precise but laborious. We therefore complemented manual segmentation analyses using machine-learning algorithms to obtain additional output data and analyze control half-domain assay experiments. All datasets were pre-processed using rotation, noise insertion, pixel intensity perturbation and translation. Cell segmentation was done on brightfield images on the half-domain assay with two machine-learning algorithms using the U-NET architecture. The first model was used to segment elongated cells and the second model to segment round-shaped cells. The first model was trained on 21 images for 50 epochs. Its accuracy was estimated at 0.97 and its recall at 0.89 (**Fig. S6**). The second model was trained on 10 images for 100 epochs and had an estimated accuracy of 0.91 and a recall of 0.73. All the training was performed using a GPU NVIDIA Quadro RTX 4000 (7.5GB RAM memory).

After identifying cells as such, round-shaped ones were assigned the category of “receivers” or “emitters” using fluorescence image information. Fluorescently labeled nuclei were also segmented using machine learning. A unique model (same for mApple tagged nuclei and for GFP-tagged nuclei) was trained (with GFP images but as well valid for RFP segmentation) over 30 epochs and its accuracy was estimated to be 0.98.

In the opto-α generated gradient (**Fig. 5**), emitters carry a mApple (red) nuclear marker, whereas receivers do not carry a nuclear marker. Cells identified as round with overlapping nuclear segmentation are identified as emitters. In the opto-Bar1-generated gradient (**Fig. 6**), emitters carry a GFP nuclear marker and were identified equivalently. Once all cells were segmented and the receivers (elongated and round shapes) are identified from the emitters, a CSV file that contains information about the identity of the segmented object (receiver or emitter), its position along the experimental axis (in µm), area (in pixels), and the mean GFP intensity in the case of fluorescent reporter receivers is produced (**Fig. 5**). For the latter, background subtraction was performed as follows: a mask was produced that is the sum of the two segmentations from the models for round and elongated shapes. The background then corresponds to the negative part of this mask in the brightfield image. To calculate the orientation of elongated objects, an ellipse was fitted using Python openCV ellipse fitting function and its angle (from 0° excluded to 180°) with respect to the x-axis was extracted (**Fig. 5F, inset**)

The data from the nuclei segmentation was used to count haploid and diploids (independently of the shape analysis, which filters cells) as a measure of cell proliferation along the experimental length in the half-domain assay (**Fig. S9**). Counting nuclei allowed us to count individual cells in regions with clumps. It also allowed the easy identification of diploid cells events in the opto-bar1 experiment (**Fig. S9**).

### 5.10 Quantification of the P*_FUS1_* transcriptional response in single cells

Quantification of the P*_FUS1_*-GFP response in the half domain assay was more precisely determined segmenting images in the GFP channel, where false positives are minimal. First the images were corrected for illumination inhomogeneities using the BaSiC package for imageJ^57^. The background profile obtained using this tool was equivalent to the background extraction done by ML but performs better in the smaller lengthscale (within-image inhomogeneities). The Cell Profiler software (Broad Institute, Cambridge, MA) was used for the quantification of the transcriptional response. The OTSU adaptive thresholding method was used for object identification in fluorescence images. Cell clumps were discarded with an object-size threshold and a form-factor filter to select rounder objects. Segmentation quality was inspected visually and empirically optimized by changing filter and threshold values. All cell segmentation data and original images are available in Zenodo.

### 5.11 Data analysis

#### Fitting pheromone gradient generated by the spatial optogenetic induction of α-factor

Due to difference in filtering, cell elongation data as well as P*_FUS1_* dependent gene-expression at single cell level had to be matched. This was done by binning the data in 100 µm wide bins and taking their mean for the downstream analysis. Fitting was done using the least squares methods on both datasets at the same time, to extract the gradient response parameters *λ* and *C* from two *θ*Z(*x*) function with different Hill coefficient *n*, normalization factor *A* and baseline *b*. Specifically, *n* was fixed to 1 for the fluorescence response, and was kept variable and equal to *n* for elongation. Only integer values between 1 and 7 were considered in the least squares minimization procedure for the *n* parameter. Fit results are summarized in **Supplementary Table 3** for different values of the Hill coefficient *n* for cell elongation (and a fixed *n* = 1 for gene expression).

## Acknowledgments

The authors would like to thank their team members for their critical reading of this manuscript. This work was mainly supported by the European Research Council grant SmartCells (724813) and received additional support from grants ANR-11-LABX-0038, ANR-10-IDEX-0001-02, ANR-16-CE12-0025-01, and the QLife Institute ANR-17-CONV-0005

## Author Contributions

AB conceived the research and contributed to all aspects of this work. MH, CE, SP, CC, and MLB contributed to optogenetic strain construction (MH, CE), experiments (MH, CE, CC), and setup of optogenetic devices (MH, SP, MLB). LC contributed image analysis. VS, YL, WJ and AA contributed the modeling of the pheromone reaction-diffusion dynamics and their fit to experimental data. AB, VS, and PH interpreted the results. AB and PH designed the study. AB, VS, YL and PH wrote the article.

## Supplementary Information

Supplementary information contains 9 figures, three tables and the mathematical model.

