## Supplement File 1 for "Optogenetic control of pheromone gradients and mating behavior in budding yeast"

### Figure S1

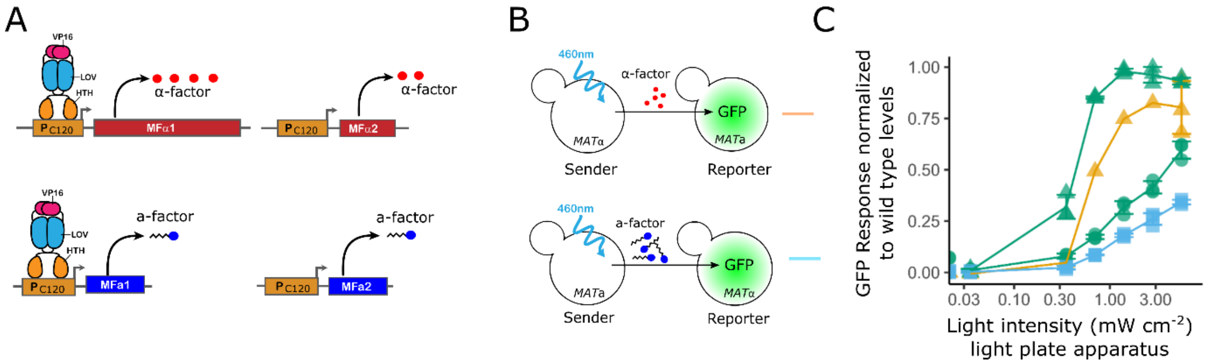


**Figure S1. Comparison of pheromone levels produced by engineered strains with native gene dosage. A.** Engineered "native" *MATα* (top) and *MAT***a** (bottom) sender strains harbor the native pheromone-producing genes with their promoters replaced with optogenetic ($P_{C120}$) ones. **B.** Schematic showing the corresponding mixes used in co-incubation assays used to quantify pheromone-mediated induction of partners (orange and blue lines correspond to traces in panel C). **C.** Gene expression responses of reporter strains in co-incubation assays measured in flow cytometry. Responses are normalized to the respective wild-type sender. The orange and blue curves correspond to the sender strains with native gene dosage (top and bottom in panel B, respectively). For comparison, the green curves corresponding to the enhanced opto-*α* strain with a wild type (circles) or Bar1Δ (triangles) *MAT***a** reporter (Fig. 2 in the main text) are shown. Notice that X-axis is logarithmic. For the orange trace, only the highest light intensity sample was replicated. Error bars are the SD of three biological replicates.

### Figure S2

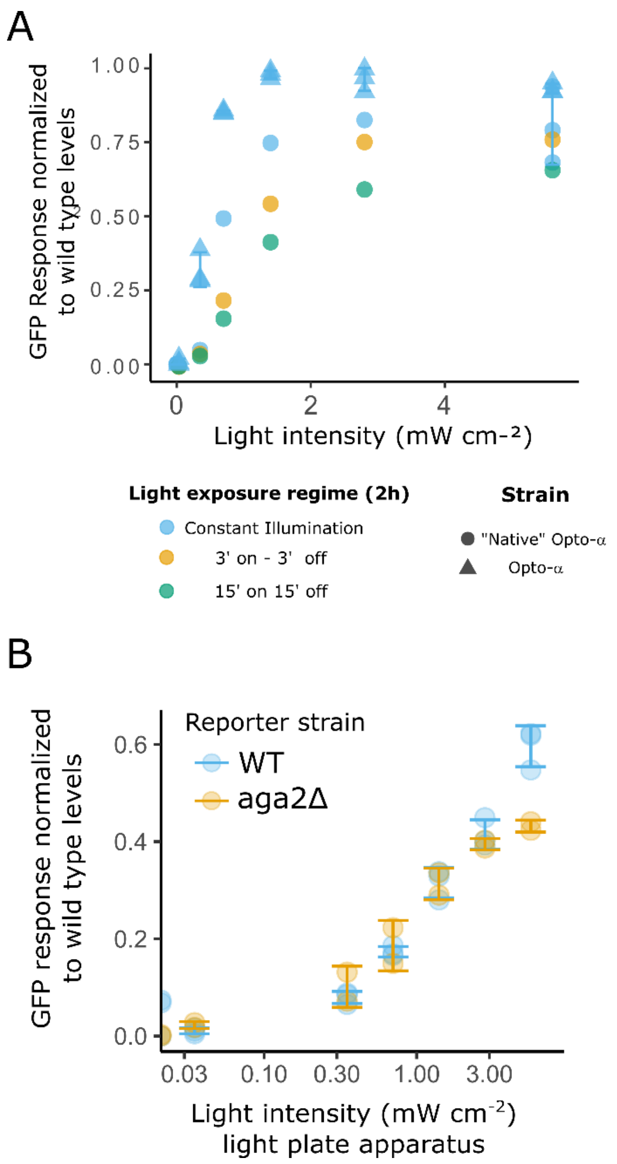


**Figure S2. Characterization of optogenetic induction. A.** Varying light exposure duty cycle in a strain without an extra-copy of MFα1 does not improve pheromone production. Co-incubation assays with different light exposure regimes. Varying duty cycles (the fraction of one period in which light is active) used on the opto-*α* strain with native gene dosage ("Native" Opto-*α*) compared to the strain with extra gene dosage (Opto-*α*). increasing the duration that cell cultures spent in the dark within a cycle, from zero to as little as three minutes reduced the overall response magnitude. **B.** Sexual agglutination does not affect light exposure. Flow cytometry results showing the $P_{FUS1}$-GFP response of a wild type *MAT***a** reporter response compared to a strain which does not form sexual aggregates. The sexual aggregation negative strain lacks the Aga2 subunit of the a-specific sexual agglutinin complex. Disaggregation before measurement was done by strong "up and down" pipetting, followed by a sonication pulse. The difference does also not arise from aggregate formation obstructing photo stimulation and/or diffusion limitation of pheromone as demonstrated by replacing the reporter strain with a non-agglutinating equivalent strain (yAA128[1]) (**Fig S2**).

### Figure S3

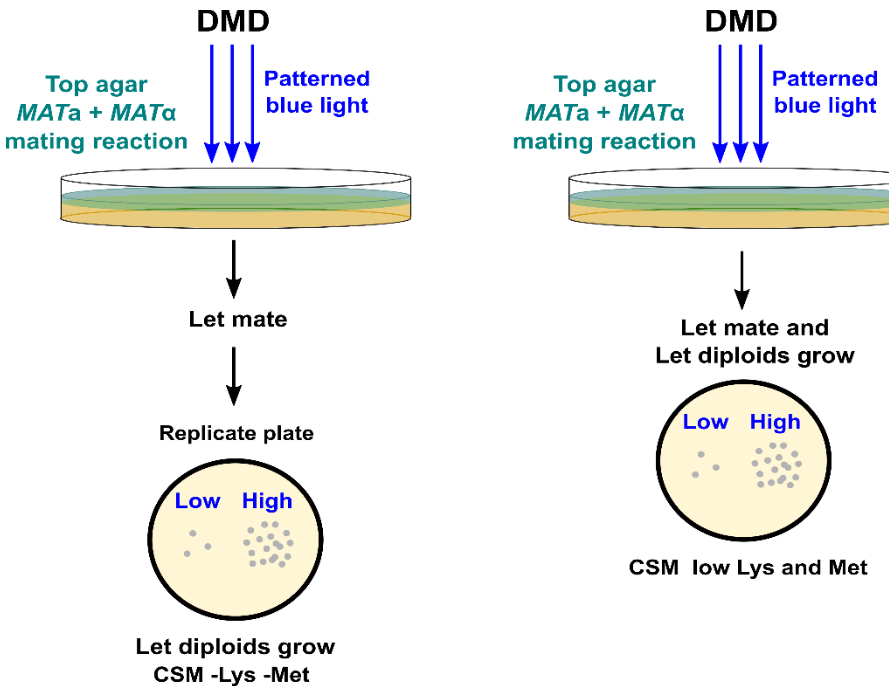


**Figure S3. Equivalent optogenetic mating assays used to quantify light-dependent mating efficiency**. Schematic for the "velvet" (left) and "direct" (right) variants of the assay used to quantify light-dependent mating efficiency. Regions of low or high light (bottom, blue) produce different diploid numbers. CSM: Complete supplement mixture. DMD: digital micromirror device. Lys: lysine. Met: Methionine (See Methods).

#### Figure S4

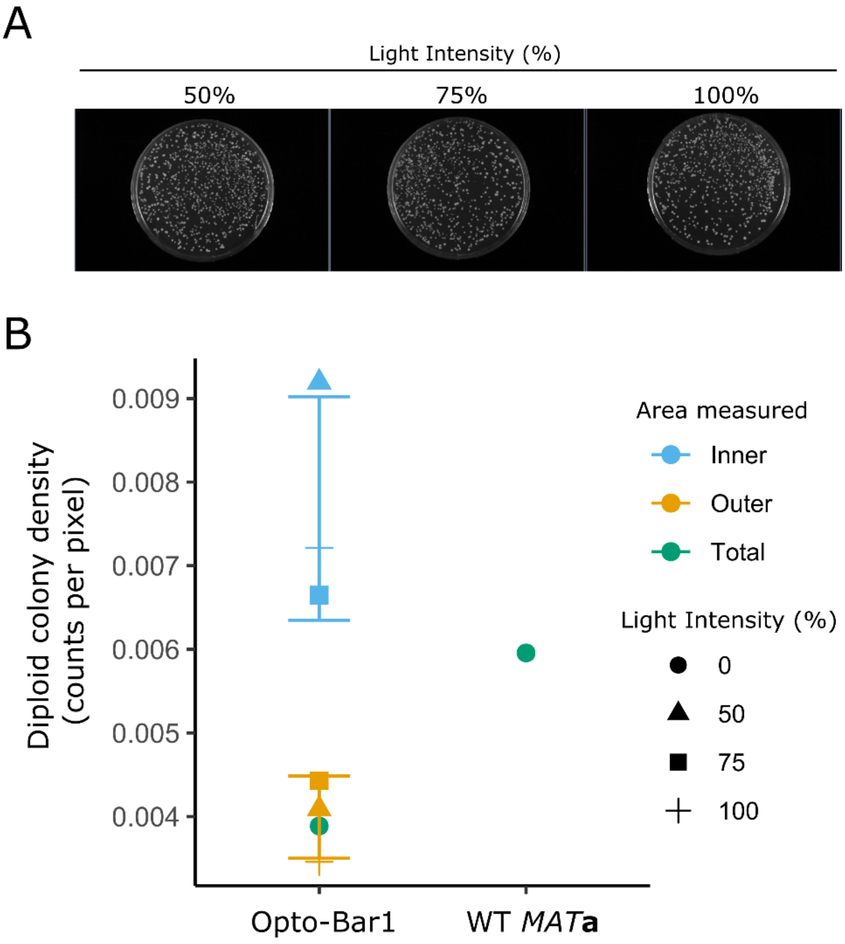


**Figure S4. Effect of optogenetically produced Bar1 on mating**. **A.** Images of plates with mating assays using an opto-Bar1/*MATα* mix Illuminated at different intensities. The leftmost image is also shown in Figure 3C (where it is the plate in the right). Images with these plates and, additionally, the "dark" and "wild type" controls in Fig. 3C were segmented to identify diploids and then counted. **B.** Quantification of diploid colony density across different illumination intensities for Illuminated and dark areas in plates (See Fig. 3) with mixes of *MATα* strain with opto-Bar1 (left). A *MAT***a**/*MATα* wild type mix is shown for comparison (right).

### Figure S5

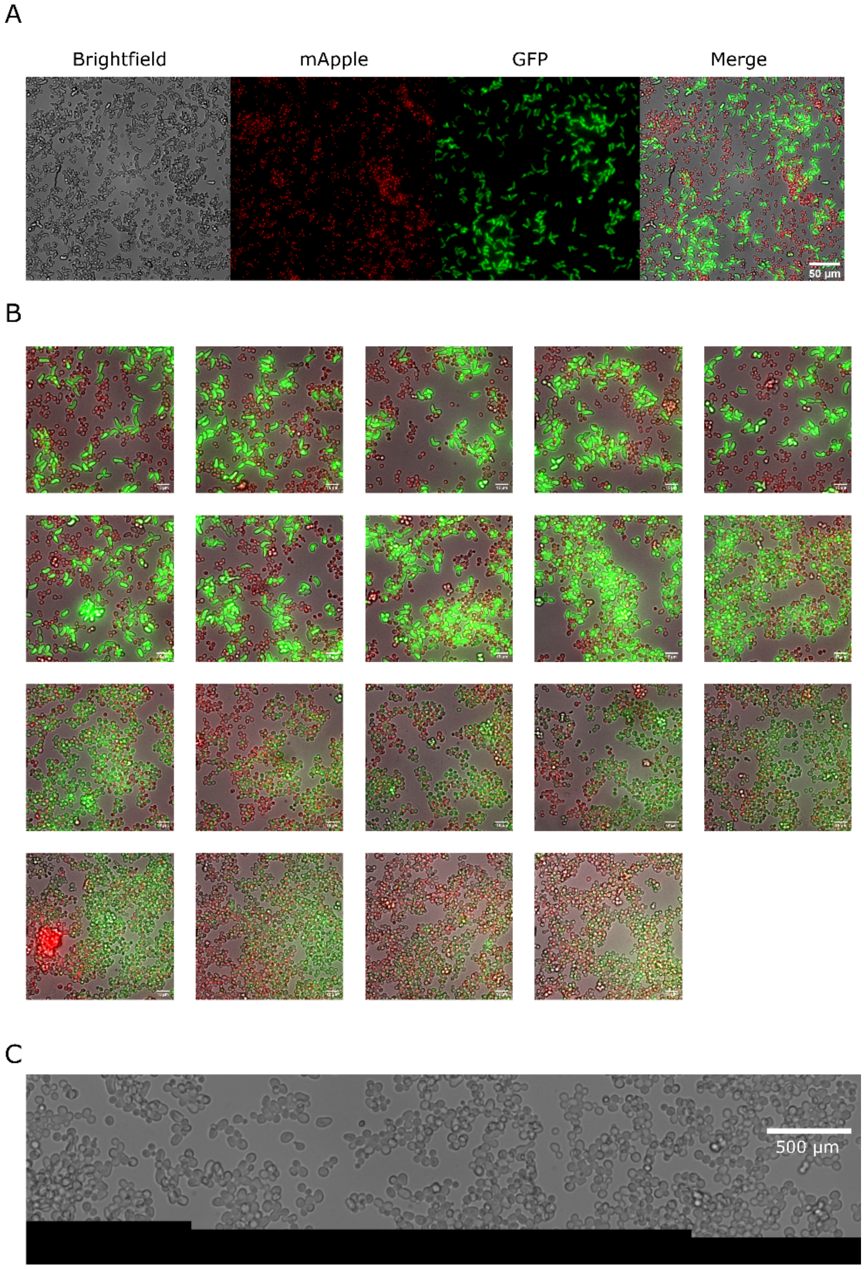


**Figure S5**. **Cell identity and overall appearance of the sampled stripe in the half-domain assay. A.** Imaging resolution and cell identity in the half-domain assay. A mixture of the opto-$\alpha$ strain (mApple, red channel) pheromone sender and the *MAT***a** Bar1Δ $\alpha$-factor receiver (GFP, green channel) in an illuminated region of the half-domain experiment showing a single tile in the stitched analyzed image. **B.** Detail of a ROI within each tile in the complete analyzed region shown from illuminated (leftmost) position to the dark (rightmost) position (read row by row from left to right.). **C.** Stitching quality. The lower end of three stitched images centered at the central image. Stitches are evidenced by the y-position realignment correction done for each image. The correction is needed because of the tilt of the 24-well plate with respect to the microscope stage frame. Scale bar: 500 µm.

### Figure S6

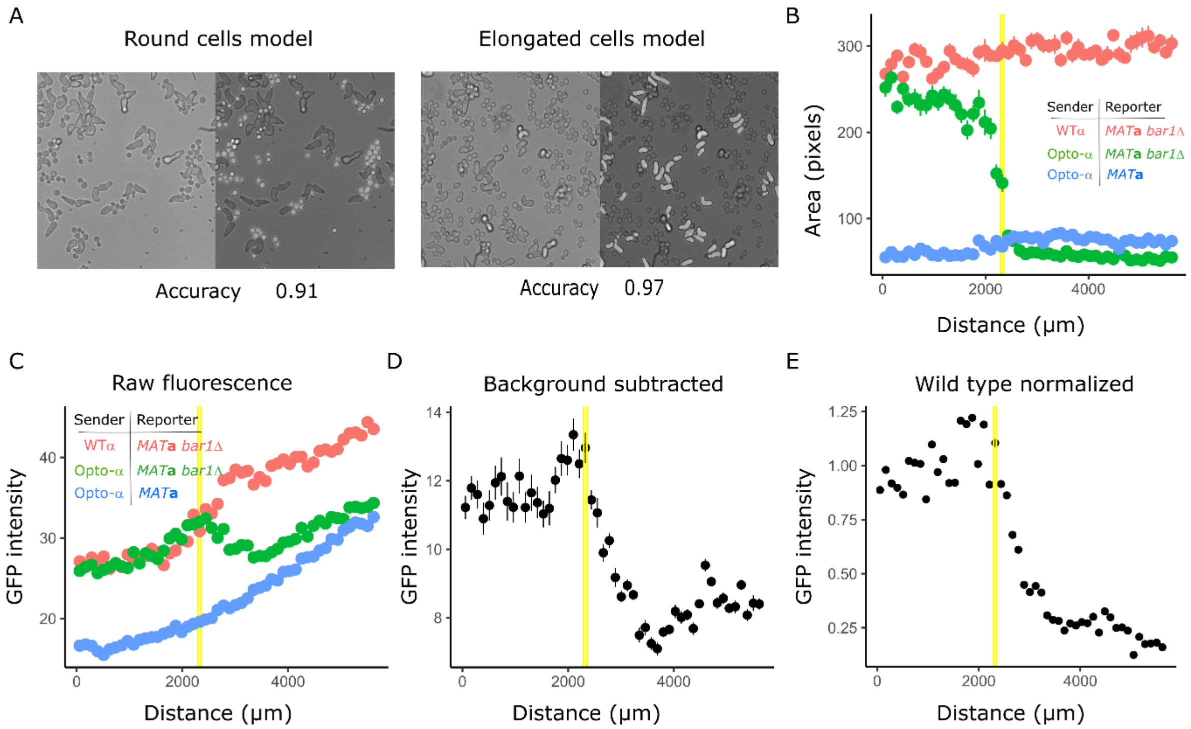


**Figure S6**. **Machine-learning based data analysis. A.** Examples of segmentations showing bright field images with their predicted masks superimposed with each one of two machine-learning models (see Methods). **B.** Segmentation was performed on three mixes. The *MAT***a**-bar1Δ receiver was incubated with Wild type MAT$\alpha$ (red), the wild-type *MAT***a** receiver was incubated with the opto-$\alpha$ strain (blue) and the *MAT***a**-bar1Δ receiver was incubated with the opto-$\alpha$ strain (green). Distance-dependency of the single-cell area of receivers as a function of distance from the measurement origin (yellow lines show the light border position of the *MAT***a**-bar1Δ / opto-$\alpha$ mix). **C-E**. Corresponding single-cell fluorescence data before normalization (C), after background subtraction (D) and after baseline (blue trace in panel C) subtraction and normalization to wild-type (red trace in panel C) levels (E).

### Figure S7

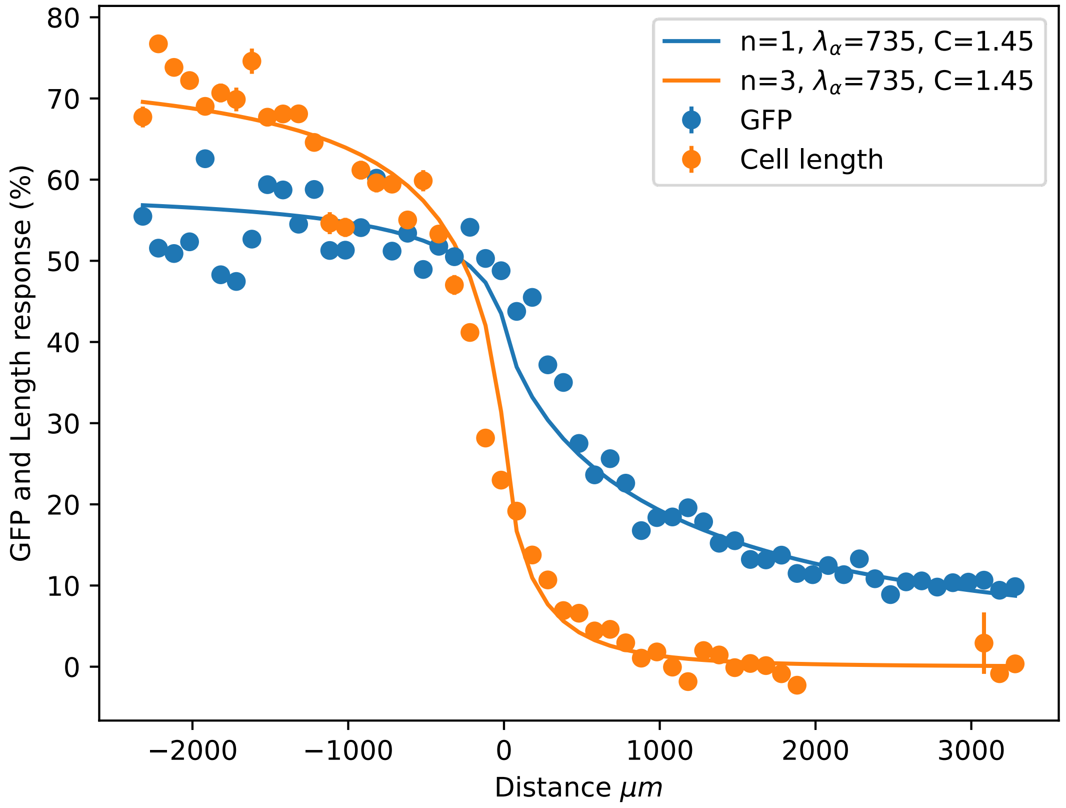


**Figure S7**. **Comparison between the morphological and gene expression responses in the half-domain experiment**. Observed gene expression from the *FUS1* promoter (GFP fluorescence) and phenotypic (cell length) responses were baseline-subtracted and renormalized, using the $\theta(x) = \left(\hat{\theta}(x) - b\right)/A$ formula and fitted values for $A$ and $b$ (Table S3), in order to obtain responses in comparable units. Points represent the mean value in each one of 53 bins, vertical lines are standard errors. Solid lines are model fits; model parameters (see main text) are shown.

#### Figure S8

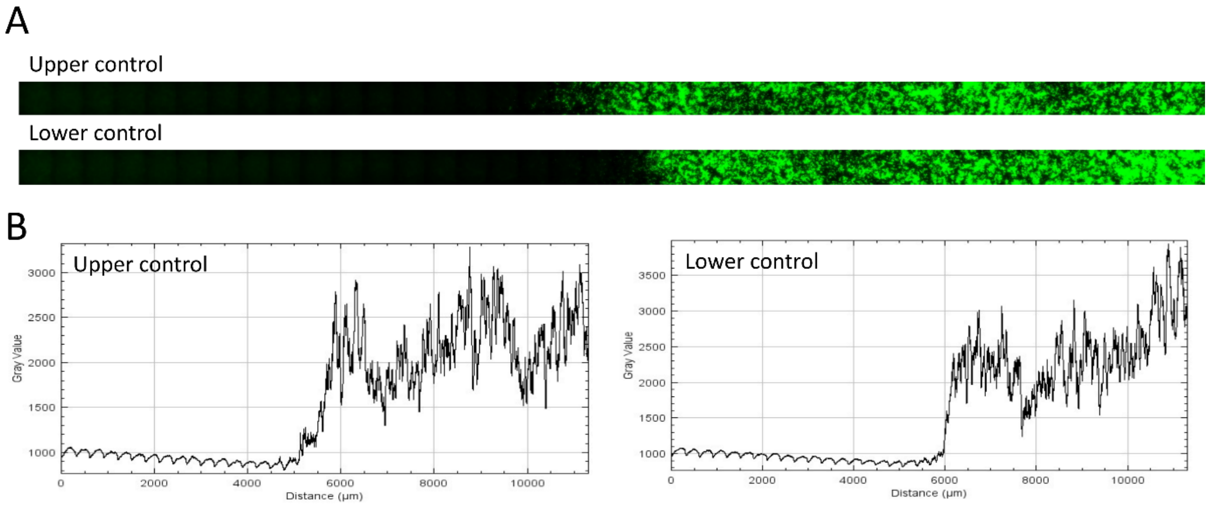


**Figure S8. Border precision in the opto-bar1 gradient generation experiment** (Fig. 6 in the main text). **A.** Stitched files for the upper and lower "biomask" light imprint controls (as done in Figure 4 in the main text). **B.** mean area intensity profile along the X-axis (correspond to the yellow vertical lines in Fig. 6 in the main text)

#### Figure S9

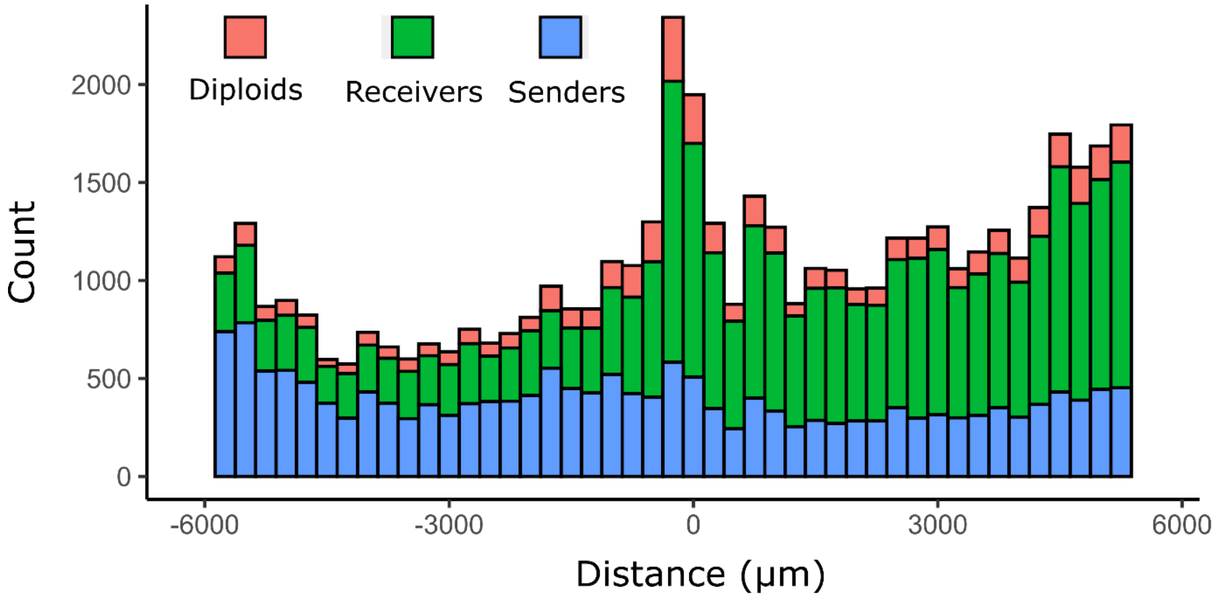


**Figure S9. Cell proliferation in the half domain assay.** Cell counts for senders of $\alpha$-factor (wild type MAT$\alpha$), receivers of alpha factor (opto-Bar1) and diploid (fusion events) in the opto-Bar1 half-domain assay (Figure 6). Bin width is 250 µm, first and last bin were removed because they contained only trace counts.

### Supplementary tables

(Starts in the next page)

| Name | Background | Mating ty | Relevant genotype | Alias / Description | Source |
|---|---|---|---|---|---|
| WT MATa | BY4741 | *MAT*a | MATa; his3D1; leu2D0; met15D0; ura3D0 | **Wild type *MAT*a** | Euroscarf |
| WT MAT$\alpha$ | BY4742 | *MAT$\alpha$* | MAT$\alpha$; his3D1; leu2D0; lys2D0; ura3D0 | **Wild type *MAT$\alpha$*** | Euroscarf |
| MH7 | BY4742 | *MAT$\alpha$* | [PC120]:MF$\alpha$1 [PC120]:MF$\alpha$2 his::EL222 | Native pheromone gene dosage opto-$\alpha$ str | This study |
| MH10 | BY4741 | *MAT*a | [PC120]:Bar1 his::EL222 | **Opto-Bar1 strain** | This study |
| MH11 | BY4742 | *MAT$\alpha$* | [PC120]:MF$\alpha$1 [PC120]:MF$\alpha$2 his::EL222 HO::PC12( MF$\alpha$1 | **Opto-$\alpha$ strain** | This study |
| MH16 | BY4742 | *MAT$\alpha$* | [PC120]:MF$\alpha$1 [PC120]:MF$\alpha$2 his::EL222 HO::PC12( MF$\alpha$1 HTB2::mApple-kanMX | **Opto-$\alpha$*** strain. An opto-$\alpha$ strain carryin mApple-HTB2 fusion as an unc fluorescent marker | This study |
| CE4 | BY4741 | *MAT*a | [PC120]:MFA1 [PC120]:MFA2 his::EL222 | Native pheromone gene dosage opto-a stra | This study |
| MH21 | BY4741 | *MAT*a | [PC120]:Bar1 his::EL222 HTB2::mApple-kanMX | An opto-Bar1strain carrying a mApple -H fusion as a nuclear fluorescent marker | This study |
| yAA198 | SEY6210a | *MAT*a | ura3::pAA35[PFUS1-Ubi(I)-sfGFP-3'FUS1URA3] aga2Δ::klTRP1 | Non-agglutinating $\alpha$-factor biosensor | Alexander Anders, MPI, Marburg |
| yAA156-1 | SEY6210 | *MAT$\alpha$* | ura3::pAA35[PFUS1-Ubi(I)-sfGFP-3'FUS1 URA3] | a-factor biosensor | Alexander Anders, MPI, Marburg |
| yAA24-1 | SEY6210a | *MAT*a | ura3::pAA35[PFUS1-Ubi(I)-sfGFP-3'FUS1 URA3] | $\alpha$-factor biosensor | Alexander Anders, MPI, Marburg |
| yAA28 | SEY6210a | *MAT*a | bar1Δ::kanMx6 ura3::pAA35[PFUS1-Ubi(I)-sfGFP-3'FUS1 URA3] | Bar1Δ $\alpha$-factor biosensor | Alexander Anders, MPI, Marburg |
| MH26 | BY4742 | *MAT$\alpha$* | WT *MAT$\alpha$* + mVenus Nuclear Tag | WT *MAT$\alpha$* carrying a mApple-HTB2 fusio a nuclear fluorescent marker | This study |

| | | | | | |
|---|---|---|---|---|---|
| yPH471 | BY4741 | *MAT*a | HO : pC120-SUC2-P2A-Venus - HIS3::(pPGK1-VP16-EL22 tCYC1) HTB2::mApple-kanMX | Biomask control for half-domain assay | Ref[2] |

**Supplementary Table 1**. *Saccharomyces cerevisiae* strains used in this study.

**Supplementary table 2.** Oligonucleotides and guide RNA sequences used in this study.

| Name | Sequence | Description |
|---|---|---|
| oPH_920 | CTAGCTCTAAAACTTCATTGACATCACTAGAGA | short guide for CRISPR insertion of PC120 upstream of MFA2 |
| oPH_921 | CTAGCTCTAAAACCTCATTAATTCATTTCTGGC | short guide for CRISPR insertion of PC120 upstream of MFA1 |
| oPH_922 | TATAGTTGTCTTTCTTTTCAGAGGATTTATCCTTCTGAGTGGCTTGTGTGGAAGCAGTGGTGATCGGTTGCATAGATTTTAGCGGCCGCG | Reverse primer for the amplification of PC120 from plasmid pYTK097 with overhangs for MFA2 promoter replacement |
| oPH_923 | GCTGTTGCATTACCACGTAATTTTGTATATAAATATCTGATAAATAACCATTTTATTTCCATCCACTTCTTTATCGCCGGGTACGTGAGT | Forward primer for the amplification of PC120 from plasmid pYTK097 with overhangs for MFA2 promoter replacement (long version) |
| oPH_924 | GCATGTATTTACCTATTCGGGAAATTTACATGACATGGATGCCATAAGGAACGAAAATGAAACATGCATGTTATCGCCGGGTACGTGAGT | Forward primer for the amplification of PC120 from plasmid pYTK097 with overhangs for MFA2 promoter replacement (short version) |
| oPH_925 | TGATAATATAGTTGTCCTTCTTTTCACTGCTGGTCTTTTCTTTTGGAGCGGCGGTAGCGGTAGATGGTTGCATAGATTTTAGCGGCCGCG | Reverse primer for the amplification of PC120 from plasmid pYTK097 with overhangs for MFA1 promoter replacement |
| oPH_926 | CCTACTGCTACGGTTGGCCCATACCTTTATTCTTTGTTCTTGTTACAAACGAGTGTGTAATTACCCAAAATTATCGCCGGGTACGTGAGT | Forward primer for the amplification of PC120 from plasmid pYTK097 with overhangs for MFA1 promoter replacement |
| oPH_927 | GATCTCTCTAGTGATGTCAATGAAGTTTTAGAGCTAG | Long guide for CRISPR insertion of PC120 upstream of MFA2 |

| | | |
|---|---|---|
| oPH_928 | GATCGCCAGAAATGAATTAATGAGGTTTTAGAGC TAG | Long guide for CRISPR insertion of PC120 upstream of MFA1 |
| oPH_775 | ATT GAG CTT CTT TTC TTG AGG AGA GAT CCA ATT TGA AGT CGG AAT AAG ATT TGC TTT CAT TAG CGT AGG CTT ATC GCC GGG TAC GTG AGT | Forward primer for the amplification of PC120 from plasmid pYTK097 with overhangs for MF(Alpha)1 promoter replacement |
| oPH_776 | TAG TGT TGA CTG GAG CAG CTA ATG CGG AGG ATG CTG CGA ATA AAA CTG CAG TAA AAA TTG AAG GAA ATC TCA TAG ATT TTA GCG GCC GCG | Reverse primer for the amplification of PC120 from plasmid pYTK097 with overhangs for MF(Alpha)1 promoter replacement |
| oPH_777 | GAT CTA GCT TCT ACT GAA AAA CAG GTT TTA GAG CTA G | Long guide for CRISPR insertion of PC120 upstream of MF(Alpha)1 |
| oPH_778 | CTA GCT CTA AAA CCT GTT TTT CAG TAG AAG CTA | short guide for CRISPR insertion of PC120 upstream of MF(Alpha)1 |
| oPH_783 | GATCTCTTTACAGCGCAGAGACGAGTTTTAGAGC TAG | Long guide for CRISPR insertion of PC120 upstream of MF(Alpha)2 |
| oPH_784 | CTAGCTCTAAAACTCGTCTCTGCGCTGTAAAGA | short guide for CRISPR insertion of PC120 upstream of MF(Alpha)2 |
| oPH_787 | TCGGGAAACTCTATAGTTTTCTGCGTTTCAGTACG CAGTTGGGCGTGCTAAAGTTGTTTTCCTAATTTGC TTATCGCCGGGTACGTGAGT | Forward primer for the amplification of PC120 from plasmid pYTK097 with overhangs for MF(Alpha)2 promoter replacement |
| oPH_788 | TATCTTCATCGGAACTAGCAGTGACAGAAACGGC CGCTAAAATAAAAGTGAGAAAGGTAGAAATGAA TTTCATAGATTTTAGCGGCCGCG | Reverse primer for the amplification of PC120 from plasmid pYTK097 with overhangs for MF(Alpha)2 promoter replacement |
| oPH_793 | GATCTCCGACATCATGCTGAAACAGTTTTAGAGC TAG | Long guide for CRISPR insertion of PC120 upstream of BAR1 |
| oPH_794 | CTAGCTCTAAAACTGTTTCAGCATGATGTCGGA | short guide for CRISPR insertion of PC120 upstream of BAR1 |

| | | |
|---|---|---|
| oPH_795 | AAAGCGCCGGTTCCTCTGACTCTAGAAGAACAA ATTGACAATGTGTCGTTGAGATACGGCAACGAGT TGGTTATCGCCGGGTACGTGAGT | Forward primer for the amplification of PC120 from plasmid pYTK097 with overhangs for Bar1 promoter replacement |
| oPH_796 | AAGCAGTAATGGTGTTAATAATCGCGAAACTCGC CAAAATAAGTTTCAAACAAAGATGATTAATTGCA GACATAGATTTTAGCGGCCGCG | Reverse primer for the amplification of PC120 from plasmid pYTK097 with overhangs for Bar1 promoter replacement |
| oPH_805 | CCA CGA AAA GTT CAC CAT AAC TTC GAA TAA AGT CGC GGA AAA AAG TAA ACA GCT ATT GCT ACT CAA ATG ATT ATC GCC GGG TAC GTG AGT | Forward primer for the amplification of the PC120-MF(ALPHA)1 cassette from genomic DNA from strain MH7 with overhangs for insertion in the HO locus |
| oPH_811 | TGG TTT TTT TCA TCC AAA ATA TTA AAT TTT ACT TTT ATT ACA TAC AAC TTT TTA AAC TAA TAT ACA CAT TGG CAT CAT AAT CAG GGA GTG | Forward primer for the amplification of the PC120-MF(ALPHA)1 cassette from genomic DNA from strain MH7 with overhangs for insertion in the HO locus |

**Supplementary Table 3**. Fit parameters for gene expression and cell elongation response functions.

**Best model fit***

| $n$ | $cost$ | $C$ ($EC_{50}$) | $\lambda_\alpha$ ($\mu m$) | $A_{gfp}$ | $A_{elo}$ | $b_{gfp}$ | $b_{elo}$ |
|---|---|---|---|---|---|---|---|
| 1 | 9660 | 0.485 | 798 | $1.19 \cdot 10^{-1}$ | 26.7 | $2.84 \cdot 10^{-3}$ | 2.74 |
| 2 | 5222 | 1.00 | 766 | $7.54 \cdot 10^{-2}$ | 17.0 | $1.67 \cdot 10^{-3}$ | 4.23 |
| **3** | **5159** | **1.45** | **735** | $\mathbf{6.28 \cdot 10^{-2}}$ | **10.7** | $\mathbf{8.63 \cdot 10^{-4}}$ | **4.55** |
| 4 | 5856 | 1.63 | 728 | $5.94 \cdot 10^{-2}$ | 8.69 | $5.10 \cdot 10^{-4}$ | 4.66 |
| 5 | 6543 | 1.64 | 715 | $5.93 \cdot 10^{-2}$ | 8.11 | $6.11 \cdot 10^{-4}$ | 4.72 |
| 6 | 7011 | 1.58 | 683 | $5.98 \cdot 10^{-2}$ | 7.92 | $1.01 \cdot 10^{-3}$ | 4.76 |
| 7 | 7291 | 1.56 | 659 | $5.99 \cdot 10^{-2}$ | 7.73 | $1.29 \cdot 10^{-3}$ | 4.77 |

* Each row presents the best fit with $n$ equal to the value in the first column, fits are obtained by optimizing the *cost* value calculated by the *scipy.optimize.least_squares* python library function at fixed $n$. The best fit for both models was chosen to minimize the *cost* across the whole column, and is presented in bold letters. Unspecified dimensions are arbitrary units.
