## Supplement File 2 for "Optogenetic control of pheromone gradients and mating behavior in budding yeast"

### The concentration distribution of $\alpha$-factors without $Bar1$

According to the experimental setup, $\alpha$-factor is secreted by $MAT\alpha$ cells (with density $\rho_\alpha$) at the $z = 0$ plane for $x < 0$ at a rate $r$. Pheromones can diffuse into the positive (right-side) half-plane and are taken up by $MATa$ cells (with density $\rho_A$) at both $x < 0$ and $x > 0$ on the $z = 0$ plane at a rate $\Omega$. Assuming translational symmetry along the $y$-axis (parallel to the interface), we can write the concentration of the $\alpha$-factor in the $x - z$ plane as follows

$$\frac{\partial c(x,z,t)}{\partial t} = D\nabla^2 c(x,z,t) + r\rho_\alpha \Theta(-x)\delta(z) - \Omega\rho_A\delta(z)c(x,z,t), \qquad (S1)$$

where $\Theta$ is the Heaviside step function and $\delta(z)$ is the Dirac delta function. At steady state, where $\frac{\partial c(x,z,t)}{\partial t} = 0$ , we have

$$\nabla^2 c(x,z) + \frac{\tilde{r}}{D}\Theta(-x)\delta(z) - \frac{\tilde{\Omega}}{D}\delta(z)c(x,z) = 0. \qquad (S2)$$

Where we rename $\tilde{r} \coloneqq r\rho_\alpha$ , $\tilde{\Omega} \coloneqq \Omega\rho_A$ . It is simple to show that $c(x,z)$ has the following symmetry

$$c(-x,z) + c(x,z) = \frac{\tilde{r}}{\tilde{\Omega}}. \qquad (S3)$$

If we substitute $\frac{\tilde{r}}{\tilde{\Omega}} - c(-x,z)$ into $(S2)$, assuming $c(x,z)$ is a solution, we find $\frac{\tilde{r}}{\tilde{\Omega}} - c(-x,z)$ is also a solution, and thus from uniqueness we can conclude $(S3)$. Specifically, we note that

$$c(0,z) = \frac{\tilde{r}}{2\tilde{\Omega}}, \qquad (S4)$$

and we also note

$$c(x,z) - \frac{\tilde{r}}{2\tilde{\Omega}} = \frac{\tilde{r}}{2\tilde{\Omega}} - c(-x,z), \qquad (S5)$$

meaning that the function $c(x,z) - \frac{\tilde{r}}{2\tilde{\Omega}}$ is odd with respect to $x$. Now, to solve $(S1)$ for $z > 0$, we have

$$\nabla^2 c(x,z) = 0, \qquad (S6)$$

and we use $(S4)$, $(S5)$ to make an informed ansatz on $c(x,z)$

$$c(x,z) = \frac{\tilde{r}}{2\tilde{\Omega}} + \sum_n \sin(k_n x)\left(A_n\, e^{k_n z} + B_n e^{-k_n z}\right). \qquad (S7)$$

We can now rewrite the production and degradation terms from $(S1)$ as a boundary condition at $z = 0$

$$\frac{\partial c(x,z)}{\partial z}|_{z=0} = -\frac{\tilde{r}}{D} + \frac{\tilde{\Omega}}{D} c(x, z=0), \tag{S8}$$

and use ($S8$) and reflecting boundary conditions on $L_x$ , $L_z$ to determine $k_n$, $A_n$ and $B_n$, and finally obtain:

$$c(x,z) = \frac{\tilde{r}}{\tilde{\Omega}} \left[ \frac{1}{2} - \frac{1}{L_x} \sum_n \frac{sin(k_n x)\, cosh\big(k_n(z - L_z)\big)}{k_n(D/\Omega\ k_n\, sinh(k_n L_z) + cosh(k_n L_z))} \right], \tag{S9}$$

with

$$k_n = \frac{\pi}{2L_x}(2n+1)\ ;\ \ n \in \mathbb{N}\,. \tag{S10}$$

In figure $S1$, we show a heat-map of ($S9$), and the corresponding flow field $-\nabla c(x,z)$. Clearly, the uptake of the $\alpha$-factor is not only due to the flow along the $z = 0$ plane; the space $z > 0$ also plays an essential role, leading to a quantitatively different result than a $1D$ model of diffusion along $z = 0$. The importance of the dimensionality on diffusion in equivalent problems has been systematically studied before[1,2].

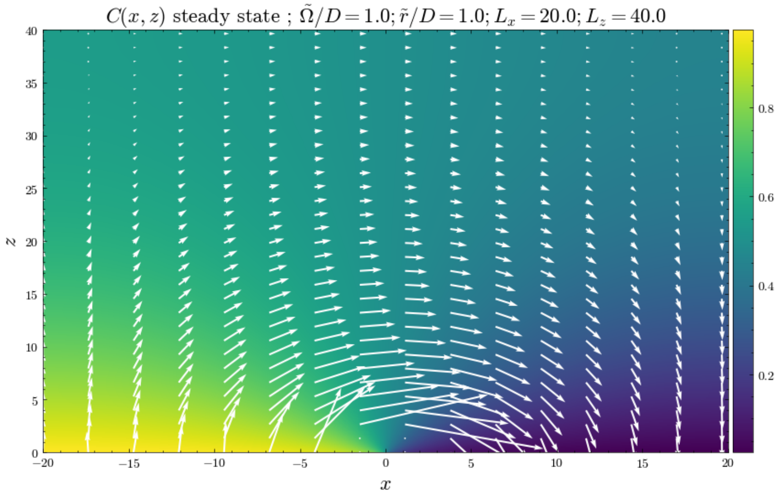


*Figure S1: Heat-map of $c(x,z)$ and the corresponding flow field. We omit a small neighborhood around $x = z = 0$ as the flow becomes very large there. Colors represent the value of $c(x,z)$.*

For $z = 0$, in the limit $k_n L_z \gg 1$ and using the well-known Fourier series expansion for square waves, we have

$$c(x,0) = \frac{\tilde{r}}{\tilde{\Omega}} \left( \Theta(-x) + \frac{1}{L_x} \sum_n \frac{sin(k_n x)}{Dk_n + \tilde{\Omega}} \right). \tag{S11}$$

We see (figure $S2$) that $(S11)$ is accurate even for $L_x$ of the order of $L_z$. We can then consider the continuous limit of $k_n$ as $L_x \to \infty$, with which we obtain

$$c(x,0) = \frac{\tilde{r}}{\tilde{\Omega}}\left[\Theta(-x) + \frac{cos\left(\frac{x}{\lambda_\alpha}\right)}{\pi} sign(x)\left(\frac{\pi}{2} - Si\left(\frac{|x|}{\lambda_\alpha}\right)\right) + \frac{sin\left(\frac{x}{\lambda_\alpha}\right)}{\pi} Ci\left(\frac{|x|}{\lambda_\alpha}\right)\right], \qquad (S12)$$

where $Si(\omega) = \int_0^\omega \frac{sin(y)}{y} dy$ and $Ci(\omega) = -\int_\omega^\infty \frac{cos(y)}{y} dy$ are the sine and cosine integrals, respectively. There is a length scale $\lambda_\alpha \equiv \frac{D}{\tilde{\Omega}}$, which is different from the case of diffusion occurring along a $1D$ $x$-axis. Asymptotically expanding $(S12)$ such that $|x| \gg \lambda_\alpha$, we obtain

$$c(x,0) = \frac{\tilde{r}}{\tilde{\Omega}}\left(\Theta(-x) + \frac{\lambda_\alpha}{\pi x}\right), \qquad (S13)$$

and we see $c(x,0)$ decays as $1/x$ in the far-field, instead of exponential decay as in the $1D$ diffusion case. In the experiment we find that $\lambda_\alpha < 1$ $(mm)$, while $L_x \sim 10$ $(mm)$, and it is evident (figure $S2$) that this is a regime where $(S13)$ is already quite accurate. When $0 < x \ll \lambda_\alpha$, we have to 1st order

$$c(x,0) = \frac{\tilde{r}}{\tilde{\Omega}\pi}\left[\frac{\pi}{2} + \frac{x}{\lambda_\alpha}\left(\ln\left(\frac{x}{\lambda_\alpha}\right) + \gamma - 1\right)\right], \qquad (S14)$$

where $\gamma$ is the Euler-Mascheroni constant. This is, again, different than the exponential $x$-dependence in the $1D$ case.

Now we turn to the microscopic picture of absorption and attempt to obtain $\lambda_\alpha$ based on the microscopic parameters. Following an established approach[3], and approximating the $MATa$ cells as independent absorbers, we can write the uptake term as $\Omega\rho_A c(x,0) = sR_A\rho_A Dfc(x,0)$, where $R_A$ is the cell radius, $f \leq 1$ is some measure of the microscopic absorption efficiency of the $MATa$ cells and $s$ is a dimensionless coefficient dependent on the effective shape one chooses to attach to the absorbers (e.g., $s = 4\pi$ in the spherical approximation). For the microscopic picture, we take $c(x,0)$ as what is typically denoted as $c_\infty$, the steady-state concentration at $z \to \infty$, assuming $c(x,z_0) \approx c(x,0)$ for some $z_0$ which is larger than all relevant microscopic length scales. This

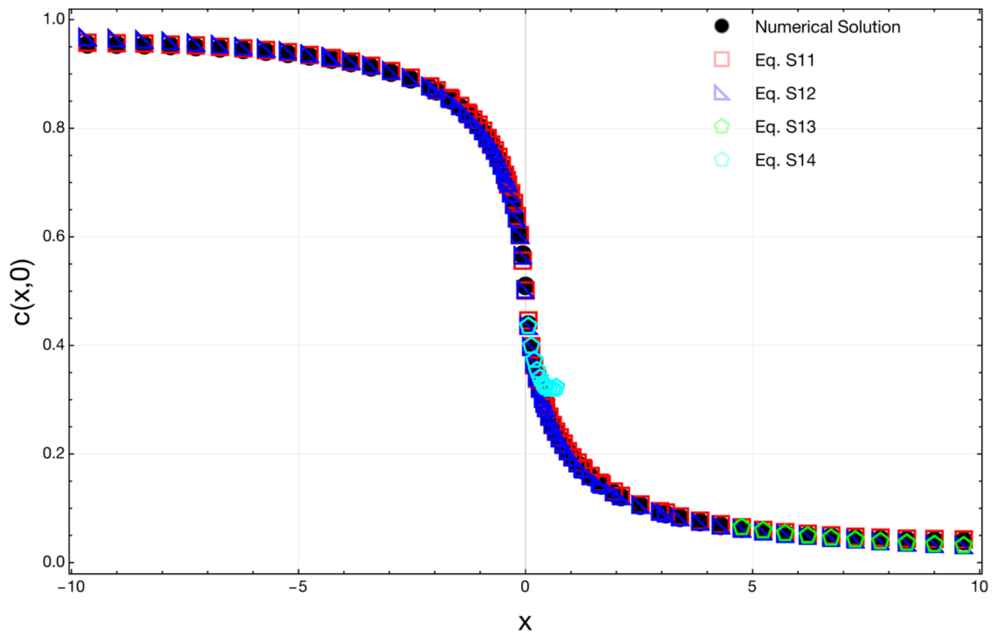


*Figure S2: Analytical and numerical solutions* $c(x, z = 0)$*, for parameter values* $\tilde{r}/D = \tilde{\Omega}/D = 1$*,* $L_x = L_z = 10$*. Numerical solution is for* $(S6)$*, based on stated boundary conditions. Far-field*$(S13)$ *and near-field*$(S14)$ *approximations are plotted in appropriate regimes for the* $x > 0$ *case.*

approximation of the absorption flux also implicitly assumes a weak spatial dependence of the steady state concentration relative to the microscopic length-scales (an assumption that breaks down as $x \to 0$). Therefore, we see that macroscopic uptake can be translated to the microscopic picture as $\Omega = sR_a Df$, and we find the length scale in term of the microscopic parameters

$$\lambda_\alpha = \frac{1}{sR_A \rho_A f}. \qquad (S15)$$

This result suggests that $\lambda_\alpha$ is independent of $D$, which is an empirically verifiable prediction. By taking the values $\lambda_\alpha = 600\mu m$, $R_A = 5\mu m$, $\rho_A = 0.01\mu m^{-2}$, we use $(S17)$ with $s = 4\pi$ to estimate $f \sim 10^{-3}$. It is, again, important to note that our approximation treats the $MATa$ cells as independent absorbers. In actuality, there are strong cooperative effects between absorbers in this problem, as seen in the famous chemoreception problem[4]. The independent absorber approximation is especially pathological for an absorbing plane (as opposed to an absorbing sphere, for instance), and one may observe that this approximation leads to certain divergences which are not physical. Cooperative effects tend to reduce the absorption relative to the independent absorber picture [4], thus one may treat these results as an upper bound for the microscopic uptake.

### Adding $Bar1$

Changing the optogenetic control scheme, we allow natural $\alpha$-factor production, and control exclusively the production of $Bar1$, confining this area of production to the $x > 0$ region and the $z = 0$ plane (see text). We also assume an effectively high activity of Bar1[5], degrading all $\alpha$-factor on the $z = 0$ plane. These are simplifications of course, but they lend themselves to a simple change in boundary conditions on $z = 0$ – for $x < 0$ we have the same boundary conditions as before, and for $x > 0$ we impose absorbing boundary conditions. These changes result in the simple solution $\tilde{c}(x,z) = \Theta(-x)\left(2c(x,z) - \frac{\tilde{r}}{\tilde{\Omega}}\right)$, $\tilde{c}(x,z)$ being the notation for the new solution with addition of $Bar1$, $c(x,z)$ being the solution for the no $Bar1$ case $(S9)$, and the rest of the parameters as previously defined. One can verify that this solution satisfies the new boundary conditions, and that it is a solution of the Laplace equation $(S6)$. Substituting $(S11)$ in the solution $\tilde{c}(x,z)$, we get

$$\tilde{c}(x,0) = \frac{\tilde{r}}{\tilde{\Omega}}\Theta(-x) \times \left(1 + \frac{2}{L_x}\sum_n \frac{sin(k_n x)}{k_n + \tilde{\Omega}/D}\right). \qquad (S16)$$

Using the previous results for the continuous approximation, we again consider the limit $L_x \to \infty$, with which we obtain

$$\tilde{c}(x,0) = \frac{\tilde{r}}{\tilde{\Omega}}\Theta(-x)\left[1 + \frac{2cos\left(\frac{x}{\lambda_\alpha}\right)}{\pi} sign(x)\left(\frac{\pi}{2} - Si\left(\frac{|x|}{\lambda_\alpha}\right)\right) + \frac{2\,sin\left(\frac{x}{\lambda_\alpha}\right)}{\pi} Ci\left(\frac{|x|}{\lambda_\alpha}\right)\right]. \qquad (S17)$$

For the $|x| \gg \lambda_\alpha$ limit we have

$$\tilde{c}(x,0) = \frac{\tilde{r}}{\tilde{\Omega}}\Theta(-x)\left(1 + \frac{2\lambda_\alpha}{\pi x}\right), \qquad (S18)$$

and for the $|x| \ll \lambda_\alpha$ limit we have

$$\tilde{c}(x,0) = \frac{\tilde{r}}{\tilde{\Omega}}\Theta(-x)\left[\frac{2x}{\pi\lambda_\alpha}\left(\ln\left(\frac{|x|}{\lambda_\alpha}\right) + \gamma - 1\right)\right]. \qquad (S18)$$

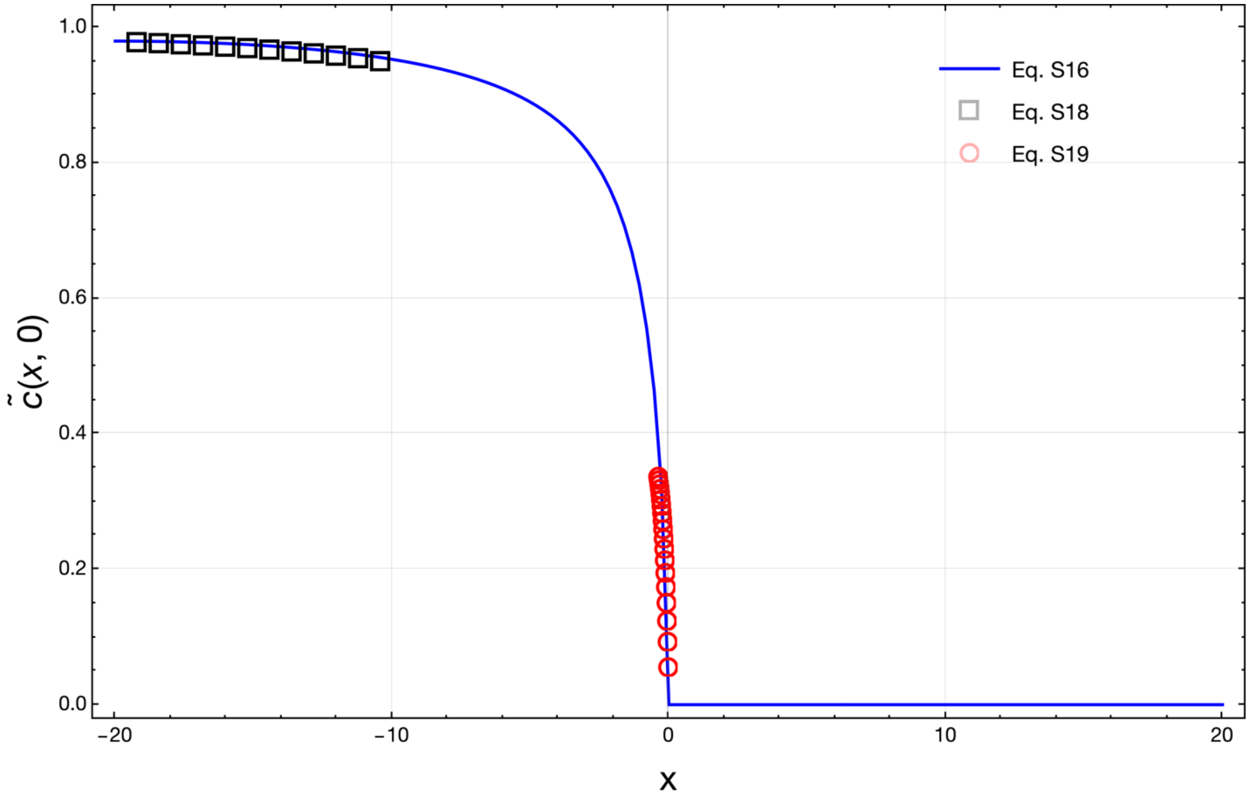


*Figure 3: Plotted solution for the concentration $\tilde{c}(x, z)$ of $\alpha$-factors on the $z = 0$ plane in the $Bar1$ case – full analytical solution $(S16)$, far-field approximation $(S18)$ and near-field approximation $(S19)$. Far-field and near-field solutions are plotted in each of the relevant regions, respectively.*
